# Systems-level analysis provides insights on methanol-based production of L-glutamate and its decarboxylation product γ-aminobutyric acid by *Bacillus methanolicus*

**DOI:** 10.1101/2024.10.14.618164

**Authors:** Marta Irla, Ingemar Nærdal, David Virant, Trygve Brautaset, Tobias Busche, Dušan Goranovič, Carsten Haupka, Stéphanie Heux, Gregor Kosec, Christian Rückert-Reed, Volker F Wendisch, Luciana F Brito, Cláudia M Vicente

## Abstract

**Background:** *Bacillus methanolicus* is the next workhorse in biotechnology using methanol, an alternative and economical one-carbon feedstock that can be obtained directly from carbon dioxide, as both carbon and energy source for the production of various value-added chemicals. The wild-type strain *B. methanolicus* MGA3 naturally overproduces l-glutamate in methanol-based fed-batch fermentations.

**Results:** Here we generated, by induced mutagenesis, an evolved *B. methanolicus* strain exhibiting enhanced l-glutamate production capability (>150%). To showcase the potential of this evolved strain, further metabolic engineering enabled the production of γ-aminobutyric acid (GABA) directly from l-glutamate, with a yield of 13.2 g/L from methanol during fed-batch fermentations. By using a systems-level analysis, encompassing whole-genome sequencing, RNA sequencing, fluxome analysis and genome-scale metabolic modelling, we were able to elucidate the metabolic and regulatory adaptations that sustain the biosynthesis of these products. The metabolism of the mutant strain evolved to prioritize energy conservation and efficient carbon utilization. Key metabolic shifts include the downregulation of energy-intensive processes such as flagellation and motility and the rerouting of carbon fluxes towards α-ketoglutarate and its derivative, l-glutamate. Moreover, we observed that transformation of the evolved strain with a GABA biosynthesis plasmid had a positive effect on l-glutamate production, likely due to an upregulation of various transaminases involved in the l-glutamate biosynthesis from α-ketoglutarate.

**Conclusions:** These results and insights provide a foundation for further rational metabolic engineering and bioprocess optimization, enhancing the industrial viability of *B. methanolicus* for sustainable production of l-glutamate and its derivatives.

## Background

Microbial bioprocesses are designed to decrease reliance on fossil-derived resources for value-added chemical production. The development of bio-based production processes is driven by the limited reserves of fossil fuels and the necessary shift towards sustainable production of important platform chemicals needed to support the growing global population. Microbial cell factories have profound value for the bio-based economy because they can selectively convert renewable feedstocks into desired products at a high yield and productivity. Such industrial workhorses include the bacterial species *Escherichia coli*, *Bacillus subtilis* and *Corynebacterium glutamicum* that are, among others, currently used for the biosynthesis of various products, including amino acids, diamines, alcohols, organic acids, enzymes and therapeutic proteins (1–4).

*Bacillus methanolicus* MGA3 possesses many features that make it a candidate to become the next industrial workhorse for sustainable bioproduction. Key traits include its natural thermophilic and methylotrophic lifestyle, and its documented high methanol consumption rate and product formation rates, including the amino acids l-glutamate and L-lysine (5). Methanol (CH3OH), a liquid and water miscible reduced derivative of carbon dioxide (CO2) that can be captured from industrial emissions, is completely utilized during microbial fermentation. During methanol-based fed-batch fermentation, *B. methanolicus* MGA3 is known to naturally produce and excrete up to 60 g/L of l-glutamate (6,7). The l-glutamate biosynthetic pathways and associated enzymes were previously characterized with a significant role of its two glutamate synthases (GLUSy), α-ketoglutarate dehydrogenase and citrate synthase in regulating l-glutamate production (8,9). l-glutamate accumulation can also be attributed to the carbon flux distribution in the central metabolism of *B. methanolicus* when grown in methanol. In this condition, methylotrophs normally do not need a complete tricarboxylic acid (TCA) cycle to fulfil their energy requirements as the reaction of methanol oxidation generates energy in the form of NADH (10). Although *B. methanolicus* is equipped with all the genes necessary for a functional TCA cycle (11–13), the levels of certain enzymes of its lower part were found to be sparse during growth on methanol (14,15). The synthesis of l-glutamate is reported to be triggered by external physiological factors including magnesium limitation (8) and the supplementation with surfactant Tween 80 (16). Finally, conversion of l-glutamate to γ-aminobutyric acid (GABA) was established in *B. methanolicus* through heterologous expression of a *Sulfobacillus thermosulfidooxidans*-derived glutamate decarboxylase gene from a GABA biosynthesis plasmid and the employment of a two-phase production strategy comprising a pH reduction from 6.5 to 4.6 (17).

In this study, we applied induced mutagenesis, screening and selection to improve l-glutamate biosynthesis without using external factors as trigger for production (18,19). l-glutamate production in selected strains was extended towards GABA biosynthesis. The resulting strains were then subjected to a comprehensive multi-omics and modelling analysis to gain a systems-level understanding of their genetic backgrounds, cell metabolism and function. Such knowledge is central in the design-build-test-learn cycle for microbial cell factory improvement while providing a rational roadmap for future work to solve bottlenecks and further boost production.

## Results and discussion

### Induced mutagenesis led to l-glutamate overproducing *B. methanolicus* strain with improved GABA production

*B. methanolicus* MGA3 mutants were obtained using classical mutagenesis combined with a high-throughput screening to identify l-glutamate overproducing clones. First, a reproducible protocol was established for growing *B. methanolicus* as single colonies on agar plates using rich complex medium supplemented with mannitol, instead of methanol, to circumvent evaporation at 50°C. In addition, a liquid pre-culture step with cells in exponential growth phase was added prior to spreading cells onto agar plates to ensure consistent cell growth on the plates.

Different mutagenesis protocols were applied and included the mutagenic agents *N*-methyl-*N*′-nitro-*N*-nitroso-guanidine (NTG), ethylmethane sulphonate (EMS) and ultraviolet (UV) light. Under these conditions, cell “kill rate” was used as a proxy measurement for DNA damage and mutagenesis rate. Parameters such as mutagenesis time, chemical mutagen concentration, UV light source distance and intensity as well as biomass input were adjusted to achieve a kill rate of <99% to avoid selection of atypical resistant variants which occur spontaneously in cell populations (20). The desired kill rate threshold was achieved for all tested mutagens. Since each mutagen causes different types of mutations (21–23), we used all three in subsequent experiments. The assay for strain mutagenesis and screening was designed as depicted in Supplementary Figure 1, Additional File 1 and described the “Materials and methods” section.

Initially, 2500 mutant clones were screened by deep-well plate cultivations, whereof roughly 30 clones exhibited a 2-fold increased l-glutamate titer compared to the parental strain *B. methanolicus* MGA3 (Figure 1A). Strain ABBM741 was selected for a second round of mutagenesis as it exhibited the most consistent improvement in l-glutamate production in different cultivation media. We repeated the mutagenesis protocol and screened an additional 800 clones for further improvements in l-glutamate production (Figure 1B).

**Figure 1.**
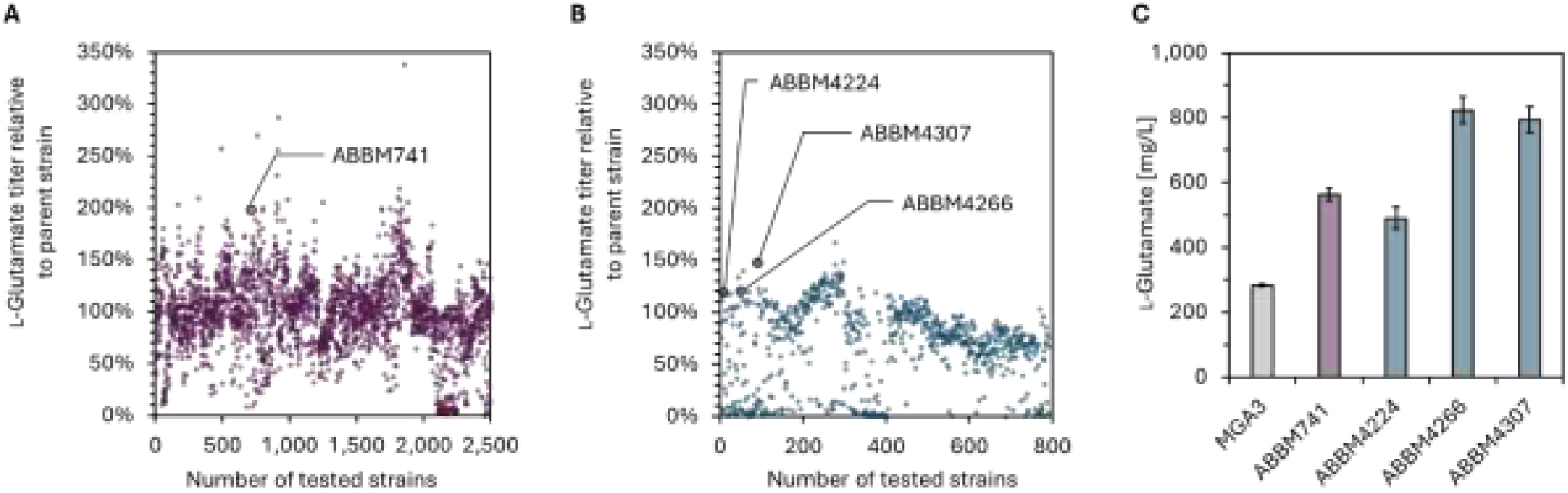
l-glutamate production during high-throughput screening and in shake flasks. A) The first generation of mutants, descending from B. methanolicus MGA3. The x axis represents the serial number of the mutant strains tested and the y axis represents the l-glutamate titer in the broth supernatant relative to the parental strain MGA3. MGA3 control cultures were present on each plate to account for variability. B) 800 clones from the second generation of mutants, descending from the top candidate ABBM741, were tested. C) l-glutamate titer in strains ABBM4224, ABBM4266 and ABBM4307 relative to their parental strains MGA3 and ABBM741.

While several strains generated in the second round of mutagenesis produced up to 50% higher l-glutamate titers than the parental strain ABBM741, only strains ABBM4224, ABBM4266 and ABBM4307 performed consistently and were further tested in biological triplicates in shake flasks instead of deep-well plates (Figure 1C). While all strains maintained increased l-glutamate production in comparison to MGA3, with strains ABBM4266 and ABBM4307 producing the highest l-glutamate titers out of the tested strains, and strain ABBM4307 was finally chosen for further tests and characterization due to its reliable growth in all experiments.

To test the capability of the evolved ABBM4307 strain to produce GABA from l-glutamate, it was transformed with the pBV2xp-*gad*^St^ plasmid which contains a *S. thermosulfidooxidans*-derived glutamate decarboxylase gene (i.e., GABA biosynthesis plasmid) to create the ABBM4307*gad* strain (17). While very low GABA titers were observed after xylose induction of *gad*^St^ expression at near neutral pH (Figure 2), significant GABA accumulation of up to 13.2 g/L was achieved for ABBM4307*gad* after culture acidification to pH 4.6. Only one third of the available l-glutamate was converted into GABA suggesting that this conversion process is a limiting step. However, the GABA titer achieved in this study increased represents a 44% increase in comparison to the GABA titer previously achieved in methanol-based fed-batch fermentations employing *B. methanolicus* MGA3(pTH1mp-*gad*^St^) (17). This is to our knowledge the highest GABA titer reported using methanol as substrate.

**Figure 2.**
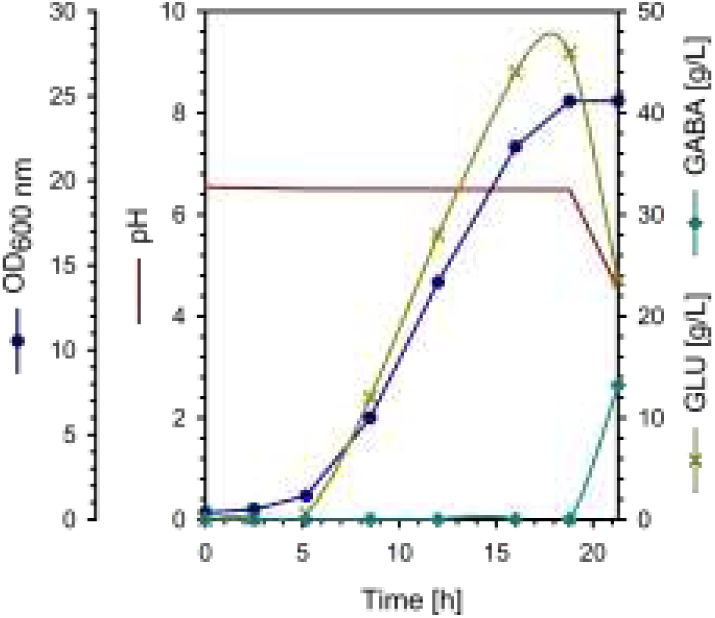
Fed-batch cultivation of strain ABBM4307gad and its methanol-based production of l-glutamate (GLU) and subsequent conversion to GABA. l-glutamate and GABA production titers were corrected for the bioreactor dilution factor due to feeding.

### Cultivation of l-glutamate and GABA overproducing strains for multi-omics characterization

To explore how overproduction of l-glutamate and its derivative GABA affect ABBM4307 and ABBM4307*gad* on different cellular levels (i.e., genome, transcriptome and fluxome), a multi-omics analysis was performed (Figure 3). The three selected strains MGA3*gad* (MGA3 transformed with pBV2xp-*gad*^St^ plasmid), ABBM4307 and ABBM4307*gad* were cultivated in a methanol-based growth medium as three biological replicates in bioreactors at controlled conditions. The multi-omics sampling was executed during the exponential growth phase, and within the optical density (OD600nm) interval of 0.6-2.2 for strain MGA3*gad* and 0.6-2.8 for stains ABBM4307 and ABBM4307*gad* (Figure 3A). During this period of approximately five hours, the biomass growth was monitored at four timepoints. The multi-omics sampling was initiated by transcriptomics sampling before and 1.5-2 h after xylose induction of *gad*^St^ expression, followed by fluxome sampling subsequent to a pulse of 3.2 g/L (100 mM) of ^13^C-labelled methanol.

**Figure 3.**
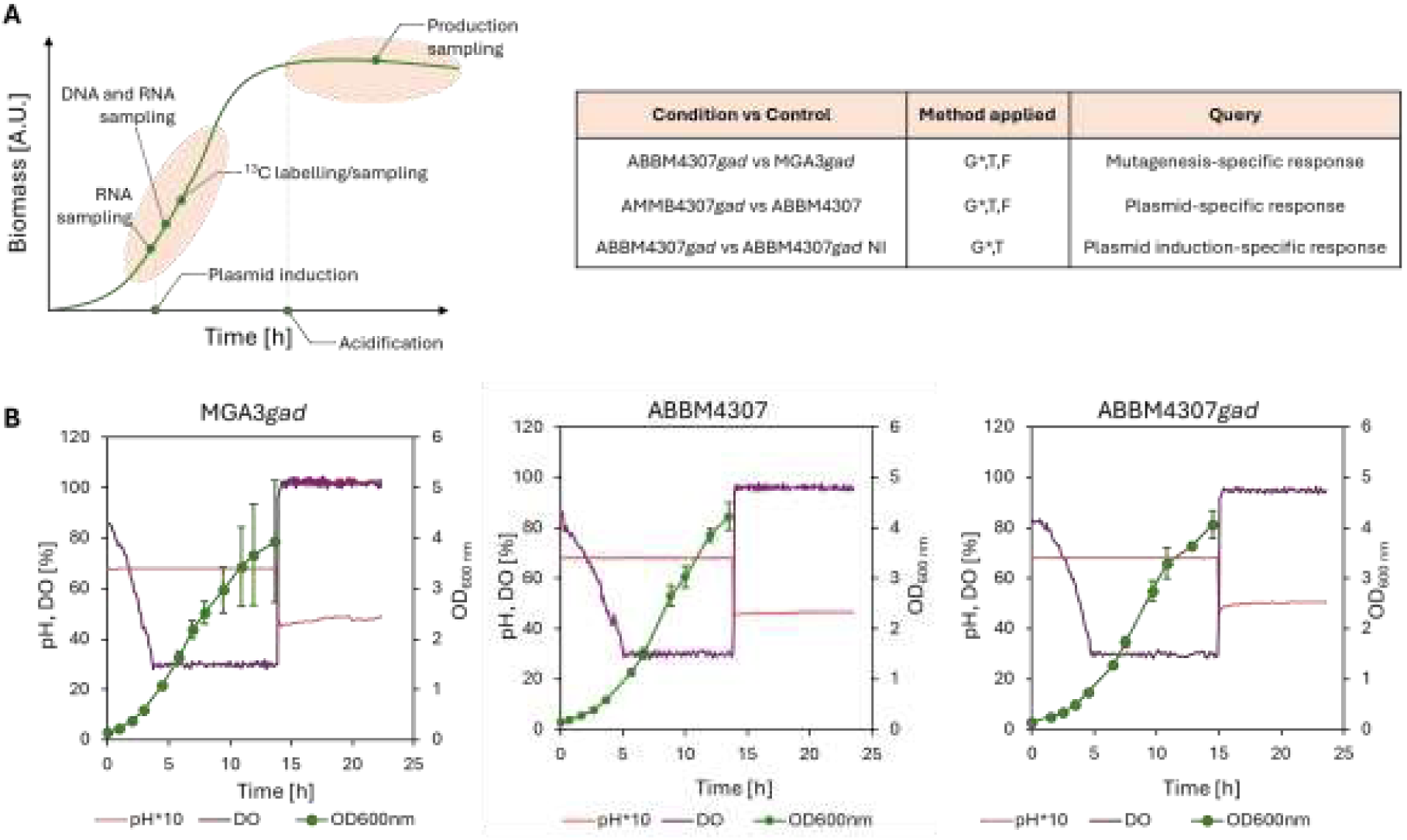
Multi-omics cultures and sampling. A) Schematic representation of the strategy of multi-omics sampling throughout bacterial growth. The table represents the control and test conditions for each query adopted in this study. G*-genomics to detect SNPs in each studied strain in comparison to the MGA3 GenBank references, T-transcriptomics, F-fluxomics. B) Plots display bacterial growth (OD600nm), pH and dissolved oxygen (DO) evolution. The growth curves are means of three biological replicates and the error bars represent their standard deviation.

The specific growth rates (µ) of the strains during the exponential phase varied and were in the range of 0.28 h^-1^-0.35 h^-1^ for all tested strains. During the multi-omics sampling phase, the triplicate cultures of all three strains displayed highly reproducible growth characteristics (Figure 3B). However, in the case of the MGA3*gad* strain, after completing the multi-omics sampling the growth of the triplicate cultures increasingly deviated with time. After the completion of the multi-omics sampling the cultures were acidified to a pH of ca 4.6 to initiate the pH-dependent conversion of produced l-glutamate to GABA by Gad^St^. Upon this pH shift the cell respiratory activity halted immediately as dissolved oxygen rapidly increased. This conversion phase lasted for nearly ten hours and samples were taken to determine the final l-glutamate and GABA titers. A separate uninoculated bioreactor, with otherwise identical conditions, was used as a control to determine the methanol evaporation rate needed for flux analysis. Sampling was performed as technical triplicates every hour. In the control bioreactor, a pulse of 3.2 g/L (100 mM) of unlabeled methanol was added at the same time when the ^13^C-methanol pulse was performed in the inoculated bioreactors.

The exometabolome samples collected during the cultivation were analyzed by NMR to assess physiological parameters such as methanol and product titers, the uptake/production rates and the evaporation constant (Table 1). The MGA3*gad* strain showed higher growth and biomass yield values, as well as methanol uptake rate, than the ABBM4307 and ABBM4307*gad* strains, in the studied conditions. As expected, the l-glutamate titer and its specific production rate were up to 5-fold higher in the evolved strains compared to parental strain. In the evolved strains, the presence of the GABA production plasmid boosted l-glutamate titer, as well as the l-glutamate production rate (Table 1). It seems that the presence of Gad^St^ creates a metabolic pull towards production of l-glutamate in ABBM4307*gad*, without a significantly increased GABA titer in comparison to MGA3*gad* (Table 1).

**Table 1.** Physiological parameters of the B. methanolicus cultures during the exponential phase and final GABA product titer. The results are presented as mean values of biological triplicates with standard deviations.

| Parameter | MGA3*gad* | ABBM4307 | ABBM4307*gad* |
| --- | --- | --- | --- |
| Biomass yield (gDCW*/g) | 0.49 ± 0.052 | 0.42 ± 0.020 | 0.43 ± 0.052 |
| Growth rate, $\mu$ (h<sup>-1</sup>) | 0.37 ± 0.01 | 0.31 ± 0.01 | 0.31 ± 0.01 |
| Substrate uptake rate (mmol/gDCW/h) | 24.0 ± 2.35 | 23.2 ± 0.92 | 22.8 ± 3.36 |
| Glutamate product titer (g/L)<sup>1</sup> | 0.03 ± 0.002 | 0.10 ± 0.025 | 0.16 ± 0.008 |
| Glutamate production rate (mmol/gDCW/h) | 0.17 ± 0.02 | 0.64 ± 0.12 | 0.81 ± 0.01 |
| Acetate product titer (g/L)<sup>1</sup> | 0.01 ± 0.004 | 0.03 ± 0.01 | 0.07 ± 0.01 |
| Acetate production rate (mmol/gDCW/h) | 0.13 ± 0.07 | 0.46 ± 0.18 | 0.89 ± 0.18 |
| Final GABA product titer (g/L)<sup>2</sup> | 0.12 ± 0.09 | ND<sup>3</sup> | 0.16 ± 0.01 |
| Evaporation constant (h<sup>-1</sup>) | 0.0471 |  |  |

\*gDCW- grams dry cell weight
<sup>1</sup>Values measured at the end of the exponential phase
<sup>2</sup>Values measured at the end of the cultivation
<sup>3</sup>Not detected

While acetate production for *B. methanolicus* during non-methylotrophic growth has only been reported once before (15), all three strains produced acetate at a different titer and rate. The evolved strain ABBM4307 produced acetate at a 3.5-fold higher rate than the MGA3*gad* strain. Here again, the introduction of the GABA production plasmid had a metabolic impact since the ABBM4307*gad* strain showed a 6.8-fold increase in acetate production rate compared to ABBM4307 (Table 1). To our knowledge, these results constitute the first observation of acetate production by *B. methanolicus* under methanol growth conditions (12,24).

### Genome and transcriptome changes cause increased l-glutamate and GABA production

To investigate the genetic basis of the induced mutagenesis and the influence of the GABA producing plasmid on l-glutamate overproduction, the genomic and the transcriptomic data of the three selected strains MGA3*gad*, ABBM4307 and ABBM4307*gad* were analyzed and compared.

First, the whole genomes from four samples, MGA3*gad*, ABBM4307, and ABBM4307*gad*, as well as ABBM4307*gad* NI (i.e., strain harvested in non-induced conditions) were sequenced. The shared and unique single nucleotide polymorphisms (SNP) identified in the genomes of analyzed strains are presented as Venn diagram in Figure 4. The analysis revealed a total of 229 SNPs across the four conditions. Notably, the mutagenesis of MGA3 to the mutant strain ABBM4307 resulted in 11 unique genetic variants within five genes encoding enzymes that belong to insertional mutagenesis process and DNA repair: recombinase (BMMGA3_02090 and BMMGA3_13340), transposase (BMMGA3_05495), and reverse transcriptase (BMMGA3_05100 and BMMGA3_06980) (Supplementary Table 1, Additional File 2). Alteration of such DNA repair systems increase mutation rates in the Gram-positive bacterium *B. subtilis* (25). Thus, ABBM4307 may have a higher potential for genetic plasticity via mutation than MGA3. This is evidenced by the fact that the genome of MGA3*gad* exhibited only a single unique SNP while ABBM4307*gad* and ABBM4307*gad* NI a total of 10 unique SNPs between both conditions indicating a 10-fold increase in mutation frequency (Figure 5; Supplementary Table 1, Additional File 2).

**Figure 4.**
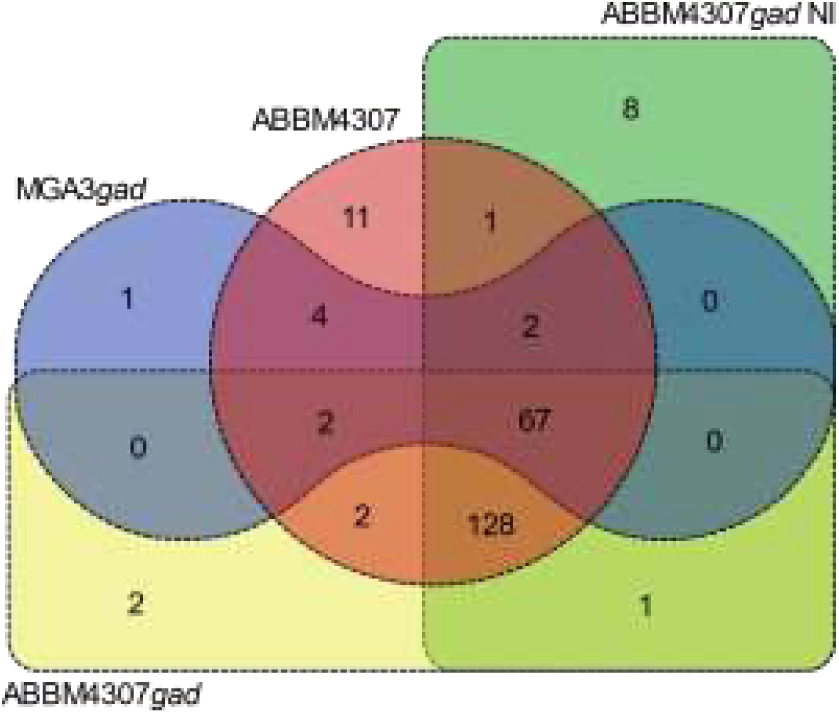
A study of genetic variants found in four l-glutamate overproduction and/or GABA production strains of B. methanolicus, MGA3gad, ABBM4307, ABBM4307gad and ABBM4307gad NI. The Venn diagram displays the shared and unique genetic variants detected in the four whole genomes.

**Figure 5.**
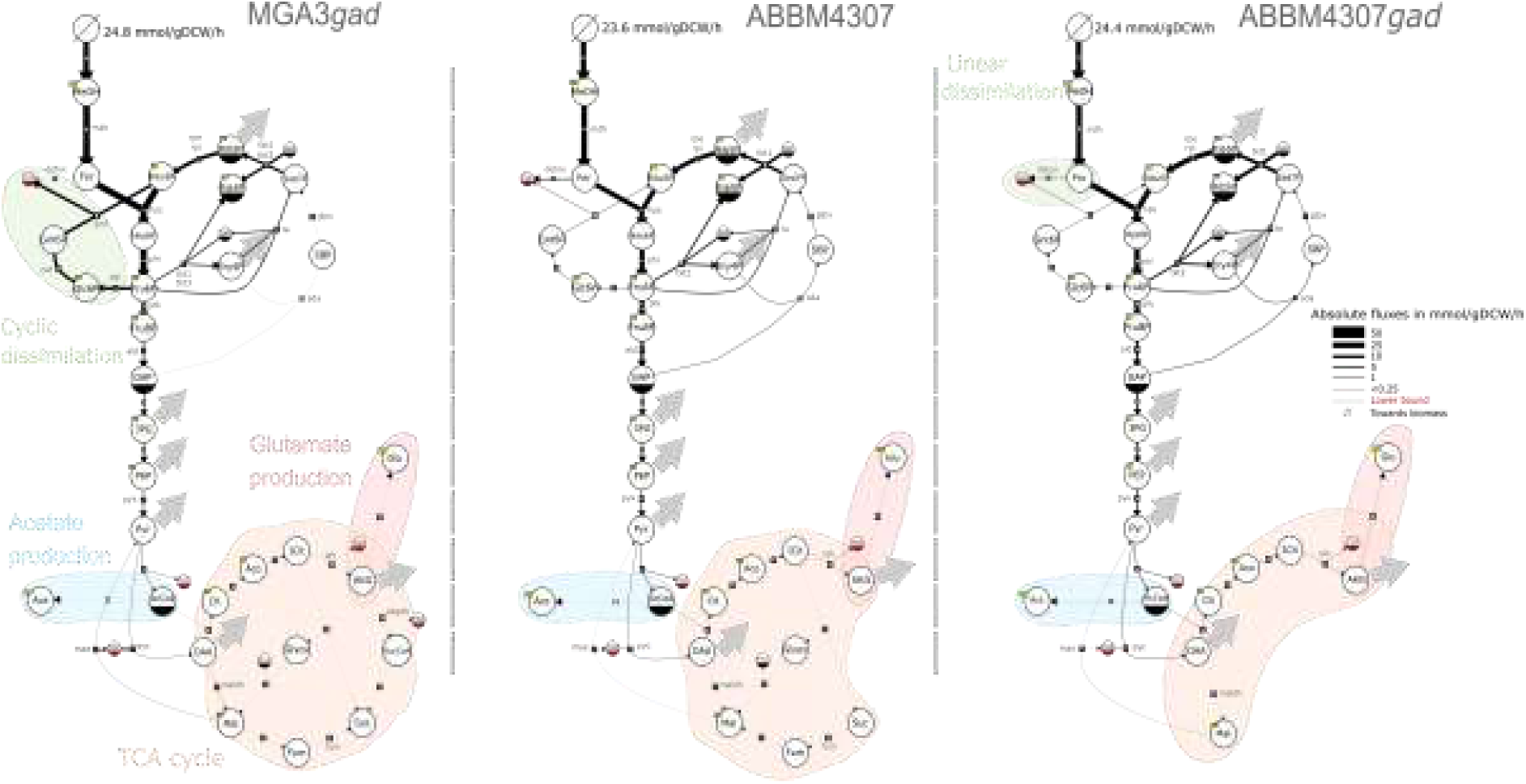
Flux maps of the central carbon metabolism illustrate the metabolic state of each strain. The thickness of the reaction lines represents the absolute flux values in mmol/gDCW/h. Flux values were averaged among three biological replicates. The direction of the reaction arrows represents the estimated net flux directionality. Large grey arrows represent a flux directed toward amino acid biosynthesis and biomass requirements for growth. Pathways discussed in the text are highlighted with a background of patch colour. Metabolites and reactions are represented following the recommendations of the Systems Biology Graphical Notation (SBGN). Metabolites are represented by circles and reactions by arrows. Metabolites which are duplicated in the same panel have a solid bottom half. Metabolites subjected to experimental measurement of ^13^C-labelling are marked with yellow boxes. Reactions towards amino acid pools and biomass are not represented. 3PG (3-phospho-D-glycerate), Ac-CoA (acetyl-CoA), Ace (acetate), Aco (aconitate), AKG (α-ketoglutarate), Cit (citrate), Ery4P (erythrose 4-phosphate), For (formaldehyde), Fru6P (fructose 6-phosphate), FruBP (fructose 1,6-bisphosphate), Fum (fumarate), GAP (glyceraldehyde 3-phosphate), Glc6P (glucose 6-phosphate), Glu (glutamate), Glyox (glyoxylate), Gnt6P (6-phosphogluconate), Hex6P (hexulose 6-phosphate), Icit (isocitrate), Mal (malate), MeOH (methanol), OAA (oxaloacetate), PEP (phosphoenolpyruvate), Pyr (pyruvate), Rib5P (ribose 5-phosphate), Ribu5P (ribulose 5-phosphate),SBP (sedoheptulose 1,7-bisphosphate), Sed7P (sedoheptulose 7-phosphate), Suc (succinate), SucCoA (succinyl-CoA), akgdh (α-ketoglutarate dehydrogenase), ald (fructose bisphosphate aldolase), detox (methanol linear detoxification), fum (fumarate reductase), glpx (sedoheptulose bisphosphatase), gnd (6-phosphogluconate dehydrogenase), hps (hexulose-phosphate synthase), idh (isocitrate dehydrogenase), mae (malic enzyme), maldh (malate synthase), mdh (methanol dehydrogenase), pfk (phosphofructokinase), pgi (glucose-6-phosphate isomerase), phi (6-phospho-3-hexuloisomerase), pyc (pyruvate carboxylase), pyk (pyruvate kinase), rpe (ribulose phosphate epimerase), rpi (ribose phosphate isomerase), sda (sedoheptulose-bisphosphate aldolase), ta (transaldolase), tkt (transketolase), zwf (glucose-6-phosphate dehydrogenase).

The unique genetic variants identified in ABBM4307*gad* NI were predominantly located in 23S rRNA genes, whereas those detected in ABBM4307*gad* were synonymous variants within hypothetical proteins (Supplementary Table 1, Additional File 2). Hence, even though transformation with the GABA biosynthesis plasmid and induction of *gad*^St^ expression in ABBM4307 resulted in unique SNPs, these variants do not appear to impact genotypic features related to l-glutamate production. The stable mutations detected in strains ABBM4307 and ABBM4307*gad*, both before and after induction, were present in different features of the metabolism of *B. methanolicus*. Key mutations were identified in genes related to flagellar assembly, amino acid biosynthesis, and fatty acid and lipid biosynthesis (Table 2).

**Table 2.** Summary of stable mutations in the genomes of the mutant strain ABBM4307. The table lists mutations across the genomes of ABBM4307, ABBM4307gad, and ABBM4307gad NI. Asterisks (*) indicate the insertion of stop codons, FS denotes frameshift insertions, and grey text represents silent mutations.

| Locus tag | Gene | Function | SNP Position | Mutation |
| --- | --- | --- | --- | --- |
| BMMGA3_01740 | *rocA* | 1-Pyrroline-5-carboxylate dehydrogenase | 322941 | Gly263Gly |
| BMMGA3_03225 | *hutH* | Histidine ammonia-lyase | 634134 | Asn119Asn |
| BMMGA3_04495 | *fabF* | 3-Oxoacyl-[acyl-carrier-protein] synthase 2 | 886520 | Ala275Thr |
| BMMGA3_04555 | *cls* | Cardiolipin synthase | 900108 | Ile247Ile |
| BMMGA3_04620 | *fabI* | Enoyl-[acyl-carrier-protein] reductase [NADH] FabI | 912024 | Arg195Lys |
| BMMGA3_05085 | *spo0A* | Histidine kinase phosphorylating Spo0A | 997567 | Arg91Lys |
| BMMGA3_05955 | *plsX* | Phosphate acyltransferase | 1168802 | Lys231Arg |
| BMMGA3_05960 | *fabD* | Malonyl CoA-acyl carrier protein transacylase | 1169371 | Ser80Ser |
| BMMGA3_06150 | *flgD* | Flagellar basal body rod modification protein | 1205541 | Thr95Ile |
| BMMGA3_06155 | *flgG* | Flagellar basal-body rod protein FlgG | 1206377 | Thr208Thr |
| BMMGA3_06395 | *dapA* | 4-Hydroxy-tetrahydrodipicolinate synthase | 1255890 | Asp243Gly |
| BMMGA3_06830 | - | Hypothetical protein | 1344179 | Gly151Gly |
| BMMGA3_08610 | *yitJ* | Bifunctional homocysteine S-methyltransferase/5,10-methylenetetrahydrofolate reductase | 1704427 | Gly407Arg |
| BMMGA3_09070 | *queE* | 7-Carboxy-7-deazaguanine synthase | 1804475 | Val242Ile |
| BMMGA3_09385 | *oxdD* | Oxalate decarboxylase OxdD | 1877514 | Gly304Arg |
| BMMGA3_09485 | *yvfT* | Sensor histidine kinase YvfT | 1895431 | Ile149Ile |
| BMMGA3_09610 | *trpC* | Indole-3-glycerol phosphate synthase | 1917486 | Gly2Glu |
| BMMGA3_09670 | *fruA* | PTS system fructose-specific EIIABC component | 1929684 | Trp498* |
| BMMGA3_09955 | *uppP4* | Undecaprenyl-diphosphatase 4 | 1985431 | Gly114Arg |
| BMMGA3_10895 | *yweB* | Cryptic catabolic NAD-specific glutamate dehydrogenase YweB | 2163086 | Cys297Tyr |
| BMMGA3_10975 | *resD* | Transcriptional regulatory protein ResD | 2180140 | Asp4FS |
| BMMGA3_11265 | *yqjD* | Putative propionyl-CoA carboxylase beta chain | 2234448 | Glu212Glu |
| BMMGA3_11330 | *bfmBC* | Dihydrolipoyl dehydrogenase | 2249633 | Gly253Asp |
| BMMGA3_12280 | *mnmA2* | tRNA-specific 2-thiouridylase MnmA 2 | 2432455 | Val167Ile |
| BMMGA3_12335 | *apt* | Adenine phosphoribosyltransferase | 2445918 | Gly139Arg |
| BMMGA3_13025 | *polA* | DNA polymerase I | 2595290 | Gly582Glu |
| BMMGA3_14735 | *hisA* | 1-(5-Phosphoribosyl)-5-[(5-phosphoribosylamino)methylideneamino]imidazole-4-carboxamide isomerase | 2913454 | Ala127Ala |
| BMMGA3_14790 | *nagA* | N-Acetylglucosamine-6-phosphate deacetylase | 2922992 | Gly67Glu |
| BMMGA3_15055 | - | Cell envelope transcriptional attenuator | 2983126 | Ala285Val |
| BMMGA3_15060 | - | Hypothetical protein | 2985037 | Ser528Leu |
| BMMGA3_16115 | *murA2* | UDP-N-acetylglucosamine 1-carboxyvinyltransferase 2 | 3213401 | Ala208Val |
| BMMGA3_16595 | *yycG* | Sensor histidine kinase YycG | 3308480 | Ser87Pro |
| BMMGA3_16635 | *purA* | Adenylosuccinate synthetase | 3313064 | Gly179Ser |

Next, a comprehensive analysis of the transcriptomic data was conducted for the following strains and conditions: MGA3*gad*, ABBM4307, ABBM4307*gad* and ABBM4307*gad* NI. Following read trimming, the total number of paired reads across all four RNA samples remained consistently at around 1.2 million reads (Supplementary Table 1, Additional File 1). The generated trimmed reads were mapped to the reference sequences of the *B. methanolicus* MGA3 chromosome and its two plasmids, pBM69 and pBM19 (respective GenBank accession numbers CP007739, CP007740, and CP007741). The alignment mapping percentages on the chromosome (∼90%), plasmids pBM19 (∼6%), and pBM69 (∼1%) remained comparable among the strains. Interestingly, when examining the GABA production plasmid in the plasmid-carrying strains, the mapping percentage on the GABA biosynthesis plasmid was 0.6% for ABBM4307*gad* NI, and significantly higher 3.2% ± 0.3% and 3.4% ± 0.1% for conditions MGA3*gad* and ABBM4307*gad*, respectively.

The transcriptomic data of the parental strain MGA3*gad* and the mutant strain ABBM4307*gad* were compared by a differential expression analysis leading to identification of a set of 421 differently expressed genes (DEGs) that were downregulated and 553 that were upregulated when comparing ABBM4307*gad* to MGA3*gad* (Supplementary Table 2, Additional File 1). We also compared whole transcriptomes ABBM4307*gad* NI to the data obtained from ABBM4307*gad* induced with 10 g/L xylose, revealing 194 upregulated and 206 downregulated DEGs in response to *gad*^St^ gene induction (Supplementary Table 3, Additional File 1). Finally, 17 DEGs were downregulated while 31 DEGs were upregulated when comparing the transcriptome of ABBM4307*gad* to ABBM4307 strain (Supplementary Table 4, Additional File 1). Overall, these results pinpoint numerous changes at both genomic and transcriptomic levels.

### Overproduction of l-glutamate affected the fluxes in the central carbohydrate network

In order to study the *in vivo* reaction rates associated with the increased l-glutamate accumulation in *B. methanolicus*, a ^13^C-metabolic flux analysis (MFA) was performed after xylose induction in the three selected strains, MGA3*gad*, ABBM4307 and ABBM4307*gad* (Figure 5).

The methanol uptake rate was not significantly different between analyzed strains (Table 1). However, the ABBM4307 and ABBM4307*gad* strains showed a slightly higher flux through the hexulose-phosphate synthase (Hps) compared to the MGA3*gad* strain (21.8 and 22.5 mmol/gDCW/h compared to 21.1 mmol/gDCW/h, respectively). As expected, a high flux toward the regeneration of the C1 acceptor pool (i.e., ribulose 5-phopshate, Ribu5P), through the non-oxidative pentose phosphate pathway (PPP) was observed in three strains, with the highest flux observed for ABBM4307*gad*. A total flux through transketolase (Tkt), fructose-bisphosphate aldolase (Sda) and transaldolase (Tal, i.e., Tala) of 17.3, 17.6 and 20.5 mmol/gDCW/h were measured, corresponding to 70, 75 and 84% of the methanol uptake rate in strains MGA3*gad*, ABBM4307 and ABBM4307*gad*, respectively (Figure 6; Supplementary Table 1, Additional File 3). The alternative route to recycle Ribu5P from fructose 6-phosphate (Fru6P) takes place through the cyclic dissimilatory ribulose monophosphate (RuMP) pathway (11,26) which operates within the oxidative PPP. The ABBM4307*gad* strain displayed a lower flux through this pathway than MGA3*gad*, with a flux through the glucose-6-phosphate dehydrogenase (Zwf) at 1.9 and 3.9 mmol/gDCW/h, respectively (Figure 5; Supplementary Table 1, Additional File 3). This coincides with the increased flux through the non-oxidative PPP in ABBM4307*gad* suggesting that in this strain Ribu5P regeneration, which does not involve carbon loss, is favored. This may have an impact on the final titer of compounds overproduced from a single-carbon substrate such as methanol. Meanwhile, regeneration of the C1 acceptor in MGA3*gad* occurred chiefly through the cyclic dissimilation pathway, likely leading to carbon loss in the form of CO2 (Figure 5).

**Figure 6.**
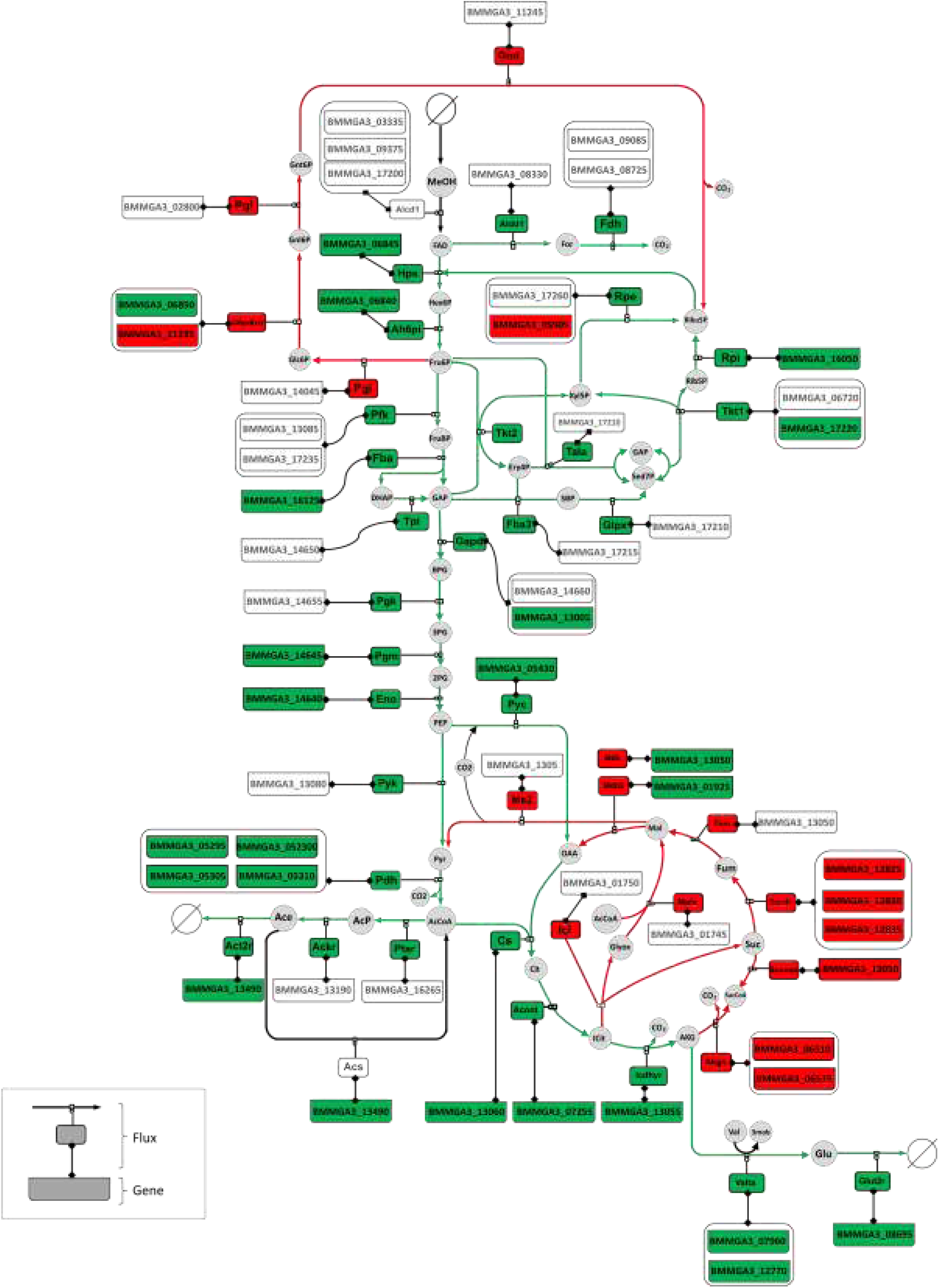
Integrative analysis of the central carbohydrate metabolism and related pathways in ABBM4307gad vs MGA3gad. Red and green indicate increased and decreased fluxes and up- and down-expressed genes, respectively. 2PG (2- phosphoglycerate), 3mob (3-methyl-2-oxobutanoate), AcP (acetyl-phosphate), BPG (1,3-bisphospho-glycerate), DHAP (dihydroxyacetone phosphate), FAD (formaldehyde), Gnl6P (6-phosphogluconolactone), Val (valine), Xyl5P (xylose 5-phosphate), Ackr (acetate kinase), Acont (aconitate hydratase), Acs (acetyl-CoA synthetase), Act2r (acetate transport), Ah6pi (hexulose-phosphate synthase), Akgs (α-ketoglutarate dehydrogenase), Alcd1 (methanol dehydrogenase), Aldd1 (formaldehyde dehydrogenase), Cs (citrate synthase), Eno (enolase), Fdh (formate dehydrogenase), G6pdh2r (glucose 6-phosphate dehydrogenase), Gapd (glyceraldehyde-3-phosphate dehydrogenase), Glut2r (l-glutamate transport), Icdhyr (isocitrate dehydrogenase), IcI (isocitrate transport), Mals (malate synthase), Mdh (malate dehydrogenase), Me2 (malic enzyme), Pgk (phosphoglycerate kinase), Pgl (6- phosphogluconolactonase), Pgm (phosphoglycerate mutase), Ptar (phosphate acetyltransferase), Sucd1 (succinate dehydrogenase), Sucoas (succinyl-CoA synthetase), Tpi (triose-phosphate isomerase), Valta (valine transaminase). Other abbreviations can be found in Figure 5.

In *B. methanolicus*, glyceraldehyde 3-phosphate (GAP) that is not used for recycling of the C1 acceptor Ribu5P is converted into pyruvate (Pyr) via a linear pathway. An increased flux through the last reaction by the pyruvate kinase (Pyk) was measured in the evolved strains (on average 4.5 and 5.4 mmol/gDCW/h in the ABBM4307 and ABBM4307*gad* strains, respectively, compared to 4.0 mmol/gDCW/h in MGA3*gad*) (Supplementary Table 1, Additional File 3). In these strains, Pyr was converted into acetyl-CoA (AcCoA) by a pyruvate dehydrogenase (Pdh), with a flux of up to 2.7 and 2.2 mmol/gDCW/h in the ABBM4307*gad* and ABBM4307 strains, respectively, compared to 1.6 mmol/gDCW/h in the MGA3*gad* strain. Interestingly, the build-up of AcCoA might have contributed to the increase of acetate production in the evolved strains with up to 3.8 and 4.8-fold more flux towards acetate production in the ABBM4307 and ABBM4307*gad* strains compared to MGA3*gad* strain, almost exactly matching the variations in the measured acetate production rates (Table 1).

Consistent with the increased flux towards Pyr and AcCoA in ABBM4307*gad*, the carbon flux through the pyruvate carboxylase (Pyc, i.e., Pc), which replenishes oxaloacetate (OAA) in the TCA cycle from Pyr, increases in the ABBM4307*gad* compared to the MGA3*gad* (2.0 mmol/gDCW/h and 1.7 mmol/gDCW/h, respectively). Unlike some methylotrophs, *B. methanolicus* carries the complete gene set for a functional TCA cycle and glyoxylate shunt (11,27). The MGA3*gad* strain showed a small flux through the TCA with an absolute flux to citrate (Cit) of 0.7 mmol/gDCW/h and very little flux afterwards, consistent with previous findings in methylotrophic conditions (12,15). Interestingly, both evolved strains, ABBM4307 and ABBM4307*gad*, presented a higher flux not only to Cit but through the TCA upper reactions from OAA to α-ketoglutarate (AKG), particularly when transformed with the GABA biosynthesis plasmid (in total, an increase of 30% and 90% in the ABBM4307 and ABBM4307*gad* strains compared with the MGA3*gad* strain, respectively). In addition, little to no flux was observed in the remaining lower reactions from AKG to malate (Mal) in these strains (Figure 5). The flux through the glyoxylate shunt was negligible, as described previously (15), with only a very low flux in the ABBM4307 strain (0.1 mmol/gDCW/h). The obtained data confirm that the TCA operates both as an oxidative and a reductive branch, with a small portion of the flux at 0.3 mmol/gDCW/h through the malate dehydrogenase (Maldh) reducing OAA to Mal in MGA3*gad*, which was even lower in the evolved strains (0.1 and 0.2 mmol/gDCW/h in the ABBM4307 and ABBM4307*gad* strains, respectively). Overall, it seems that in the evolved strains metabolic flux is directed towards AKG supporting the high production of its derivative, l-glutamate. Indeed, the flux towards l-glutamate production was 3.6-fold and 6.9-fold higher in the ABBM4307 and ABBM4307*gad* strains, respectively.

### Integration of fluxome with genome and transcriptome data revealed strain-specific altered regulatory landscape

In order to identify the potential regulatory mechanisms involved in the increased l-glutamate accumulation in the evolved strain transformed with the GABA biosynthesis plasmid, genome wide flux data predicted with a genome scale model (GSM) of *B. methanolicus* and fluxes measured with the MFA were integrated with the genome and transcriptome data for strains MGA3*gad* and ABBM4307*gad*. Since a GSM of *B. methanolicus* was not available, it was constructed in the course of this study as described in the “Materials and methods” section. A strong correlation between simulated (predicted with the GSM) and estimated (measured by MFA) fluxes was obtained (Supplementary Figure 2, Additional File 1) demonstrating a high accuracy of the GSM. Next, the measured (Supplementary Table 2, Additional File 3) and simulated (Supplementary Table 3, Additional File 3) flux distributions were combined and used to calculate the log2 fold change values for each of the 1020 reactions belonging to the GSM. This dataset was combined with the table containing the list of 974 genes, which were significantly (*p*-value < 0.01) and differentially (log2 fold change value outside the range [-0.25; 0.25]) expressed in ABBM4307*gad* in comparison to MGA3*gad* (Supplementary Table 4, Additional File 3) and the table containing the 27 genes which showed a stable (and not silent) mutation in ABBM4307*gad* in comparison to MGA3*gad* (Table 2). This facilitated the classification of adverse and similar responses of a specific gene at the genomic, transcriptional and flux levels (Supplementary Table 4, Additional File 3). Out of the 1020 reactions, 14 were associated with a mutated gene (Supplementary Table 5, Additional File 3). This includes: (i) one reaction associated with only differential change at the flux level (log2 fold change value outside the range [-0.25; 0.25]); (ii) four reactions associated with only changes at the transcript level and (iii) nine reactions with no changes at the flux and transcript level. Of the 1020 reactions, 691 were not associated with any differential change, either at gene level or at flux level (Supplementary Table 6, Additional File 3). The rest of the dataset was composed of: (i) 26 reactions with changes both in the flux and expression indicating that the regulation of these reactions is occurring at a transcriptional level (i.e., transcriptional regulation) (Supplementary Table 7, Additional File 3); (ii) 34 reactions with changes in flux but not in expression (Supplementary Table 8, Additional File 3); and (iii) 269 reactions with changes in expression but not in the flux (Supplementary Table 9, Additional File 3). In the two latter cases, it is suggested that the regulation of these reactions occurs at a level other than transcription (i.e., post-transcriptional regulation). This type of regulation includes modifications of the enzymatic activity by phosphorylation or acetylation (i.e., post-translation modification) and/or by the binding of inhibitors or activator (i.e., allosteric regulation). Such modifications can increase or decrease enzyme activity, thus modifying flux in the absence of changes in gene expression, or conversely counterbalance changes in enzyme levels due to gene expression, thus preventing any change in flux. Regardless of the regulation type, all these metabolic reactions play an important role in the l-glutamate and GABA overproduction which is further described in the following chapters.

### The central carbohydrate network is tightly regulated in the ABBM4307*gad* strain harboring the GABA biosynthesis plasmid

Notably, all the reactions of the central carbohydrate metabolism were regulated at both transcriptomic and post-transcriptomic level (Figure 6). While the fluxes of all the reactions involved in the formaldehyde assimilation were increased in ABBM4307*gad* compared to MGA3*gad*, only four of them (Hps, Ah6pi (i.e., Phi), Tkt, Fba and Rpi) showed increased fluxes along with an increase of the transcript levels of their respective genes. This is consistent with the upregulation of the *hxlR* gene encoding a transcriptional regulator which controls the expression of the chromosomal genes *hps* (BMMGA3_06845) and *phi* (BMMGA3_06840) (28). Surprisingly, the flux through the Rpe reaction was increased despite the downregulation of one of its encoding genes (Figure 6). Metabolic reactions involved in the cyclic dissimilation of the RuMP showed changes in fluxes in ABBM4307*gad* compared to MGA3*gad* but were unaffected at the gene expression levels, except for G6pdh2r (i.e., Zwf) which is encoded by two alternative genes in the genome of *B. methanolicus*. The BMMGA3_06850 gene is co-localized with the *hps*-*phi* operon in the chromosome and was upregulated in ABBM4307*gad*, while BMMGA3_11235 is located elsewhere on the chromosome and was downregulated in ABBM4307*gad* compared to MGA3*gad* (11). It is not known which one of the two genes is responsible for the Zwf activity. However, the increase of the flux through Zwf corresponding to the upregulation of BMMGA3_11235 seems to indicate a role of this gene in the regulation of the G6pdh2r-catalyzed reaction.

Interestingly, the regulation of reactions involved in the conversions of GAP into Pyr is governed by both transcriptional and post-transcriptional mechanisms. The conversion of Pyr into AcCoA by Pdh showed coordinated changes between the flux and the transcription of its encoding operon (i.e., BMMGA3_05295-BMMGA3_05305). Our data highlights transcriptional regulation of Pdh, as well as almost all reactions of the TCA cycle (Supplementary Table 2, Additional File 2). The coordinated response of transcript abundances and fluxes in the TCA cycle observed in the evolved strain compared to the parental strain seem to be adapted to promote AKG and hence l-glutamate accumulation.

In *B. methanolicus*, l-glutamate synthesis has been described to occur from AKG through the activity a glutamate dehydrogenase (GLUDxi) encoded by BMMGA3_10895 and two glutamate synthases (GLUSy) that are encoded by BMMGA3_01940-BMMGA3_01945 and BMMGA3_03480 (8). In the present study, predicted fluxes through these reactions were null and transcripts levels were similar in all tested strains for the 3 genes encoding for GLUSy. In addition, the gene encoding GLUDxi was downregulated which is consistent with the identified SNP (890G>A), resulting in the missense mutation Cys297Tyr (Table 2), which based on an InterProScan analysis is in an active site of the enzyme. Our GSM analysis identified a valine transaminase (Valta) putatively encoded by BMMGA3_07960 and BMMGA3_12770 as responsible for 85% of the l-glutamate production in both ABBM4307*gad* and ABBM4307. Interestingly, this reaction showed increased fluxes along with the upregulation of both encoding genes in ABBM4307*gad* compared to MGA3*gad*. In *Lactococcus lactis* subsp. lactis NCDO 2118, l-glutamate biosynthesis from AKG does not involve glutamate dehydrogenase and exhibits low-level activity of GLUSy and glutamine synthetase (29). At the same time, it demonstrates high transaminase activity, functioning in the opposite direction of the typical branched-chain amino acid biosynthetic process. This transaminase converts AKG to l-glutamate, utilizing a range of amino acids as nitrogen donors, with a particular preference for isoleucine, leucine, and valine (29).

In contrast to the reactions involved in the l-glutamate production which are transcriptionally regulated, the ones involved in production of acetate i.e., acetate kinase AckR (encoded by BMMGA3_13190) and phosphate acetyltransferase Ptar (encoded by BMMGA3_16265), were post-transcriptionally regulated with the exception of the transporter Act2r (encoded by BMMGA3_13490). The metabolic capacities of these enzymes were not limited by the levels of their mRNAs since ABBM4307*gad* in comparison to MGA3*gad* was able to direct an up to 52 log2 fold change increase in flux through these enzymes without a differential change in transcript level. Notably, a similar type of regulation has been recently demonstrated for acetate production in *E. coli* (30).

Overall, these findings underscore the coordinated regulation of gene expression encoding enzymes of the central carbohydrate metabolism, which appears to be an important process governing the observed metabolic behavior. From an evolutionary perspective, such coordinated changes can only be explained by mutations affecting global regulators involved in the transcriptional control of many genes. According to the transcriptomic data, 26 transcriptional regulators are differentially expressed between the ABBM4307*gad* and MGA3*gad* strains (Supplementary Table 2, Additional File 2). In addition, genomic data showed a mutation in the transcriptional regulatory protein ResD (Table 2). In *B. subtilis*, ResD activates transcription of diverse genes in response to oxygen limitation (31), however its role in *B. methanolicus* has not been described yet.

### Post-transcriptional regulation affects reactions beyond central carbohydrate metabolic network in the ABBM4307*gad* strain

More than 90% of the post-transcriptionally regulated reactions are outside of the set of core biochemical reactions involved in the central carbohydrate metabolism (Table 3). Interestingly, several reactions of the same metabolic pathway were coregulated. The metabolic pathways with the most pronounced coregulated metabolic reactions include the biosynthesis of cell wall components, amino acids, vitamins, fatty acids, nucleotides, cofactors, coenzymes and prosthetic groups (Table 3).

**Table 3.**
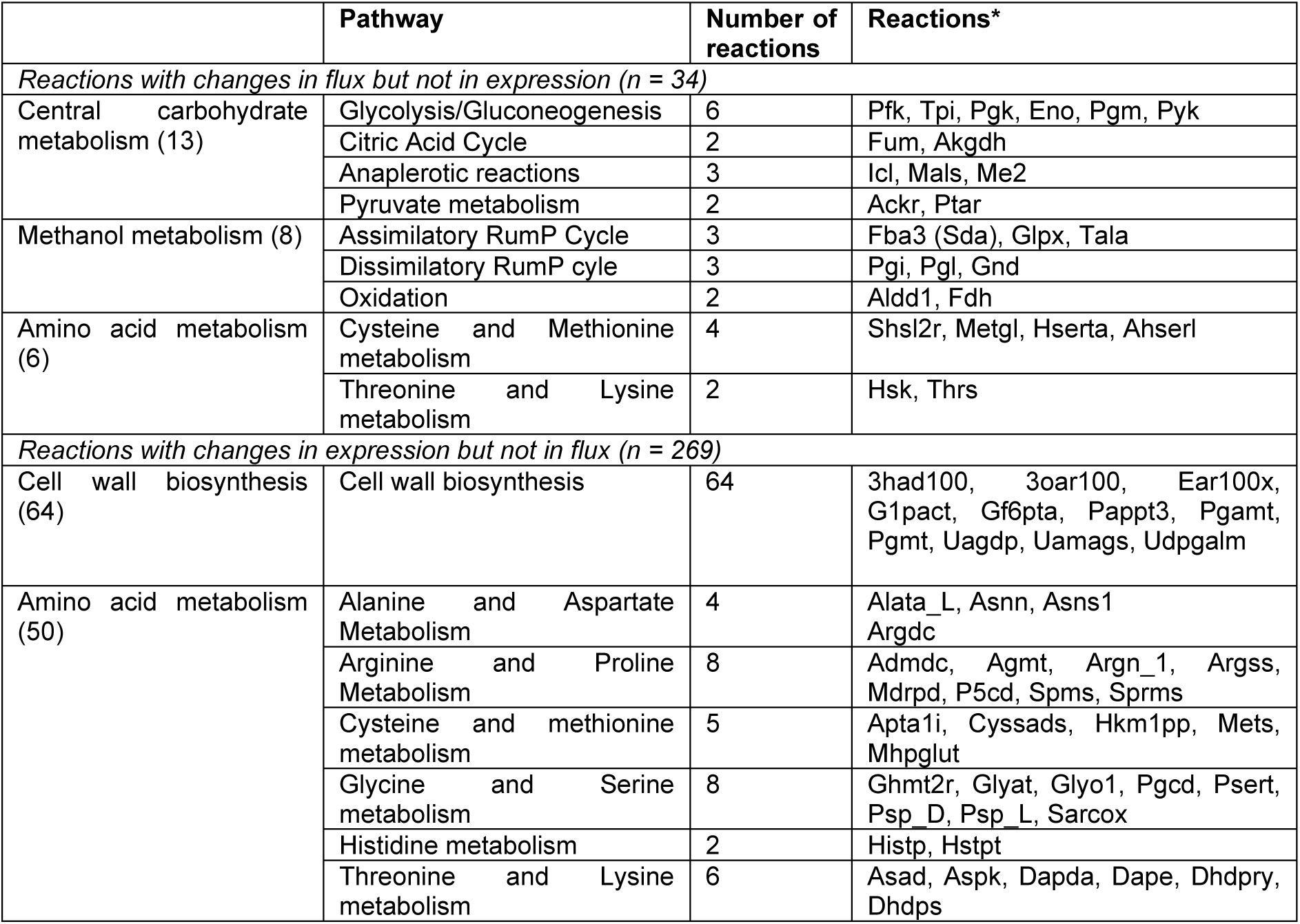
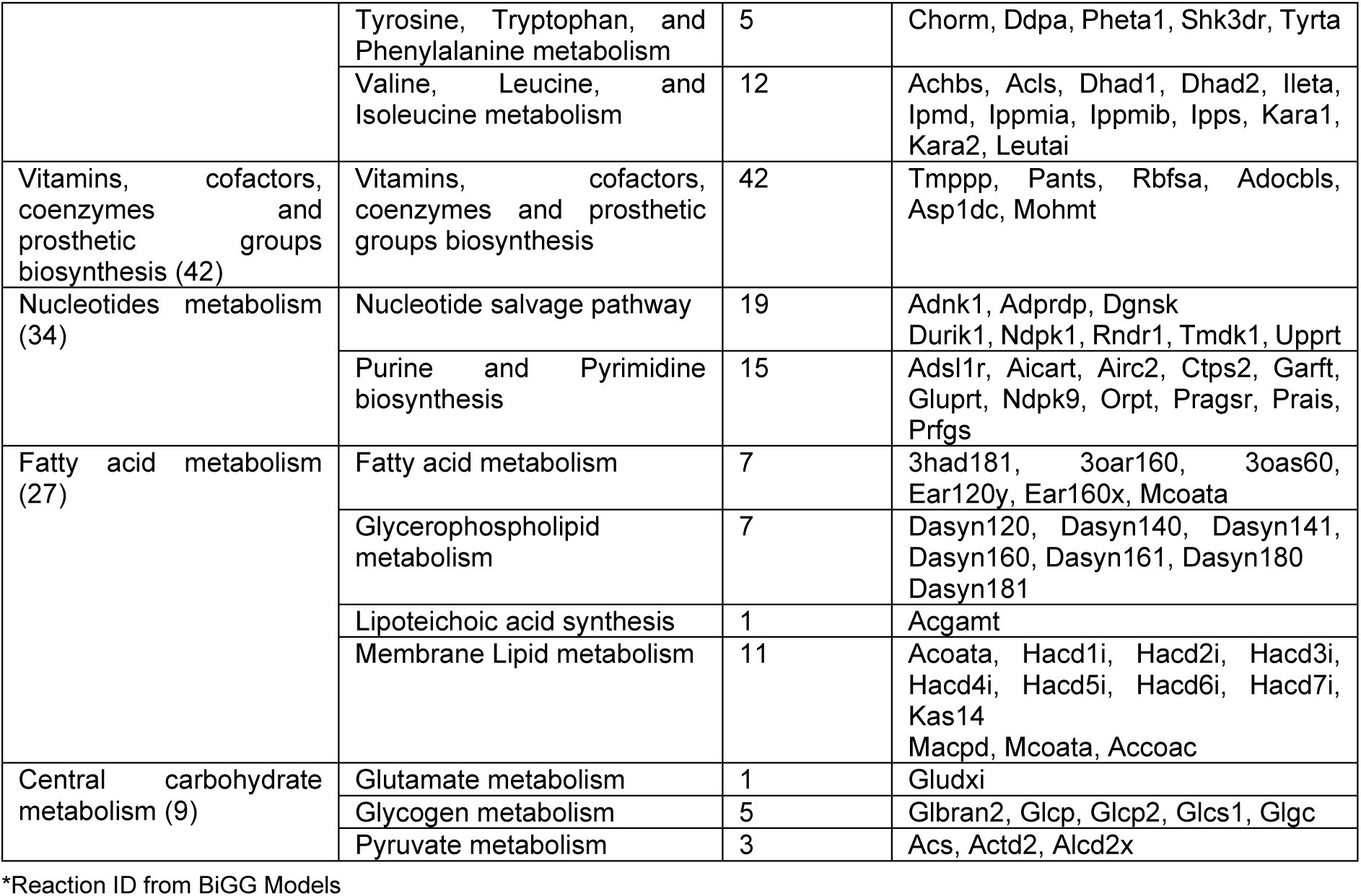
Functional classification of post-transcriptionally regulated reactions in strain ABBM4307gad compared to strain MGA3gad. The KEGG and BioCyc pathways were used for grouping.

Our results demonstrated that the 64 reactions involved in the teichoic acid, nucleotide sugars and peptidoglycan metabolism, which are crucial components of cell wall in Gram-positive bacteria, were post-transcriptionally regulated in the mutant strain ABBM4307*gad* vs MGA3*gad* (32). Interestingly, more than 94% of the reactions showed an increased level of transcripts in the corresponding encoding genes indicating preference towards cell wall preservation in ABBM4307*gad*. However, none of the reactions associated with mutated genes showed changes in the expressions of the encoding genes (Table 2; Supplementary Table 9, Additional File 3).

Proteins constitute about half of the dry biomass of *B. methanolicus*, and expression of amino acid biosynthesis genes is tightly regulated to meet that demand during growth (15). l-glutamate may play a critical role in this process since all nitrogen from ammonium is fixed by glutamate dehydrogenase and delivered to other branched-chain amino acid by transamination. Therefore, its overproduction is expected to strongly impact the biosynthesis of the other 19 amino acids. This is indeed the case since 50 metabolic reactions involved in the synthesis of all amino acids showed changes in expression of their encoding genes but not in the fluxes, while six metabolic reactions showed changes in the flux but not in the expression of their encoding genes (Table 3). Interestingly, the leucine, isoleucine and valine biosynthesis pathways genes in ABBM4307*gad* strain were upregulated in comparison to the parental strain MGA3*gad* (Supplementary Table 2, Additional File 2). It is worth noting that valine, isoleucine, and leucine are converted by the aminotransferases into α-ketoisovalerate, α-keto-β-methylvalerate, and α-ketoisocaproate, respectively, through the condensation with AKG. These metabolic reactions result in the production of l-glutamate, and pyridoxine is suggested to serve as a potential cofactor in these processes. As mentioned before, the genes BMMGA3_07960 and BMMGA3_12770 encoding the valine transaminase Valta and involved in the overproduction of l-glutamate in ABBM4307*gad* were upregulated in ABBM4307*gad* in comparison to MGA3*gad* but not in ABBM4307*gad* when compared to ABBM4307 (Supplementary Table 2-3, Additional File 2). In addition, the enzyme encoding gene involved in another transaminase reaction (BMMGA3_05040), which is putatively involved in the conversion of 4-methylthio-2-oxobutanoate and l-glutamate to AKG as well as L-methionine biosynthesis, was downregulated in ABBM4307*gad* compared to ABMA4707 (Figure 6; Supplementary Table 3, Additional File 2). This could explain why ABBM4307*gad* produces more l-glutamate than ABBM4307. Catabolism of leucine, isoleucine and valine leads to pantothenate and coenzyme A (CoA) formation, and the genes encoding the corresponding enzymes for pantothenate biosynthetic reactions (Asp1dc and Mohmt encoded by BMMGA3_10610 and BMMGA3_10620, respectively) were upregulated in ABBM4307*gad* compared to MGA3*gad* (Table 3). Mutations occurred in three genes encoding the enzymes in the reactions of amino acid metabolism (BMMGA3_06395, BMMGA3_09610, BMMGA3_11265) but without consequence either on the flux or on the expression (Supplementary Table 4, Additional File 3) except for BMMGA3_06395 which was upregulated in ABBM4307*gad* compared to MGA3*gad*. This is consistent with the fact that these mutations do not occur in the active site according to an InterProScan analysis.

Various reactions involved in the biosynthesis of vitamins, cofactors, coenzymes and prosthetic groups showed changes in the expression of their encoding genes, but not at flux level. In particular, genes BMMGA3_01785, BMMGA3_01095, BMMGA3_03010, BMMGA3_09890, BMMGA3_10615, BMMGA3_01205 encoding the enzymes in reactions Tmppp, Rbfsa, Pants, Adocbls, which are involved in the biosynthesis of B-group vitamins (B1, B2, B5, B12), were upregulated in the strain ABBM4307*gad* compared to MGA3*gad*.

*B. methanolicus* uses an energy-efficient salvage pathway for vitamin B12 biosynthesis, involving the import of exogenous corrinoids through ATP-binding cassette (ABC) transporters and consistently, two gene encoding ABC transporter permeases (BMMGA3_01175, BMMGA3_09460) were upregulated (Supplementary Table 2, Additional File 2) (33). Vitamins B1, B2, and B5 serve as essential cofactors in reactions of the carbohydrate metabolism (e.g., Pdh, Tkt) and CoA biosynthesis. Interestingly, genes BMMGA3_16300, BMMGA3_04410, BMMGA3_01115, encoding the enzymes in the reactions involved in molybdenum cofactor (Moco) synthesis were upregulated in ABBM4307*gad* (Supplementary Table 2, Additional File 2). While these reactions are absent in the GSM, it is interesting to notice that Moco is required for the functioning of the transcriptionally regulated reaction of formaldehyde dehydrogenase (Fdh) involved in the detoxification of formaldehyde (Figure 6) (34). The upregulation of various vitamin biosynthetic clusters is consistent with the increased need for cofactors of the up-regulated reactions involved in central carbohydrate metabolism and various biosynthetic pathways.

Purines and pyrimidines are the building blocks for the synthesis of nucleotides. They are also needed to produce riboflavin and flavoproteins like flavin mononucleotide and flavin adenine dinucleotide, which are important cofactors used in the electron transfer chain in bacteria (35,36). Several reactions involved in the purine biosynthetic pathway showed increased levels of transcripts in the corresponding enzymes’ encoding genes (Supplementary Table 2, Additional File 2). Among all the reactions involved in nucleotide metabolism only a few showed a stronger increase in the genes transcript’s levels that encode the corresponding enzyme in comparison to others. For instance, the four reactions (Rndr1-4) which catalyse the conversion nucleoside 5’-diphosphates to deoxynucleoside 5’-diphosphate (a basic component of DNA) showed a 4 log2 fold change in their enzymes encoded gene’s expression (BMMGA3_09290 and BMMGA3_09295) in ABBMG407*gad* in comparison to MGA3*gad*. This typically indicates that DNA replication is increased in ABBMG407*gad* although the fluxes through those reactions remain stable. One mutation occurred in one gene involved in a reaction of purine and pyrimidine metabolism but without consequence either on the flux or on the expression (Supplementary Table 5, Additional File 3).

Changes in the expression of genes involved in reactions of fatty acid metabolism were visible in the ABBM4307*gad* strain. In particular, enzymes involved in glycerophospholipid metabolism showed a downregulation in the encoding genes while the genes encoding membrane lipid and fatty acid metabolism showed an upregulation. Fatty acids are key building blocks for the phospholipid components of cell membranes, and most of the reactions involved in the initiation and elongation of fatty acids (Table 3) showed increased levels of the corresponding gene transcripts (i.e., *fabDFGHILZ* genes, Supplementary Table 2, Additional File 2). In addition, two reactions (EAR160x and 3OAS60) had mutations in their enzyme encoding genes, BMMGA3_04620 and BMMGA3_04490, BMMGA3_04495, concurrent with upregulation of one of them (Supplementary Table 4, Additional File 3). Glycerophospholipids are the main components of the lipid bilayer that constitutes cell membranes, and they contribute to the fluidity and flexibility of the membrane. These changes in membrane biosynthesis could affect l-glutamate export, similarly to mechanisms observed in *C. glutamicum* where membrane alterations influence the export through mechanosensitive transmembrane channel proteins activated by the force-from-lipids (37).

Our data demonstrates that post-transcriptional regulation of the reactions in ABBMG407*gad* compared to MGA3*gad* mainly occurs at the transcriptomic level, while affecting various parts of the metabolism. From an evolutionary perspective, a change in gene expression without a corresponding change in metabolic flux can be explained by several mechanisms that reflect the robustness and adaptability of the biological systems. It is possible that the genomic mutations generated via induced mutagenesis result in many transcriptional changes that are buffered due to redundancy in the system (e.g., multiple enzymes catalyzing the same reaction) and regulatory mechanisms (i.e., post-translational modifications or allosteric regulations) ensuring steady fluxes even when there are changes in gene expression. It is conceivable that laboratorial evolution has also favored phenotypic plasticity in the ABBMG407*gad* strain, enabling it to regulate certain genes (e.g., transaminases) in anticipation of increased metabolic activity (e.g., production of l-glutamate to sustain GABA synthesis), but without immediately increasing flux. This enables the cell to rapidly increase its activity when needed, without wasting resources when it does not.

### Important cellular processes are affected by l-glutamate overproduction

As described above, l-glutamate overproduction in *B. methanolicus* significantly alters carbon flux, affecting energy balance and the electron transport chain (ETC) in the ABBM4307*gad* strain. Specifically, the coordinated downregulation of the expression level and metabolic flux of succinate dehydrogenase (Sucd1, Figure 6) observed in the ABBM4307*gad* strain compared to MGA3*gad* might reduce electron transfer to ubiquinone, which is converted to ubiquinol in the ETC. Additionally, reduced flux through the TCA cycle decreases NADH formation, limiting the electron supply to complex I of the ETC. These changes impact the expression of genes encoding ETC components (Supplementary Figure 3, Additional File 1; Supplementary Table 2, Additional File 2), leading to a broad effect on oxidative phosphorylation. The ETC of *B. methanolicus* has not been described in detail yet but the inspection of genomic sequence suggests similarities with the ETC of *B. subtilis*. The ETC in *B. methanolicus* includes NADH dehydrogenases NDH-I, encoded by the *nuo* locus and NDH-II encoded by either *yutJ* or *yumB,* and branches ending in either quinol or cytochrome c terminal oxidases similarly to *B. subtilis* (38,39). Despite the overall downregulation of ETC-related genes, the *cydABCD* gene cluster, encoding cytochrome *bd* quinol oxidase and a heterodimeric ATP-binding cassette transporter, was upregulated (Supplementary Table 2, Additional File 2) suggesting a compensatory mechanism under low oxygen conditions as it has been observed in *E. coli* (40).

The transcriptomic data indicates that l-glutamate overproduction correlates with decreased synthesis and activity of flagella together with the downregulation of chemotaxis genes and upregulation of genes related with sporulation and biofilm formation. Genes involved in flagella structure and function (e.g., *flgD*, *fli*, *flg* and *flh)* were downregulated ABBM4307*gad* compared with MGA3*gad* (Supplementary Table 2, Additional File 2), and a mutation in the *flgD* gene was identified in ABBM4307 (Table 2). It is worth noticing that the transcriptomic response of the flagella-related processes persisted in the evolved strain expressing the GABA biosynthesis plasmid both induced and uninduced (Supplementary Table 3-4, Additional File 2). Chemotaxis genes, including a cluster coding for the chemotaxis response regulator protein-glutamate methylesterase (*cheB*) and chemotaxis central regulator (*cheA*), were also downregulated. In addition, the *spo0A* gene, a master regulator of sporulation and biofilm formation in *B. methanolicus*, was upregulated in ABBM4307*gad* compared to MGA3*gad* (Supplementary Table 2, Additional File 2), enhancing sporulation while downregulating biofilm formation (41). These changes in gene expression reflect a strategic shift to minimize energy intensive process (i.e., flagella assembly and chemotaxis) (42,43) and carbon intensive process (i.e., biofilm formation) (41), reallocating resources towards l-glutamate production.

## Conclusions

By engineering *B. methanolicus* MGA3, we significantly enhanced its l-glutamate production through induced mutagenesis and subsequently extended the metabolic pathway to synthesize GABA at high titers. System-level analyses combining genomic, transcriptomic, fluxomic and modelling revealed that regulation of various central carbohydrate reactions at the transcriptional level plays an important role in adjusting fluxes from methanol to l-glutamate, thus ensuring a high titer of its derivative GABA. However, post-transcriptional reactions in different parts of the metabolism are equally important to ensure readiness while optimizing resource use. As outlined earlier, the integration of various omics-level approaches has yielded detailed quantitative data for biological network analysis in *B. methanolicus*, serving as a crucial prerequisite for targeted strain improvement. It would be beneficial to incorporate proteome analysis in future studies and expand metabolome measurements to include intermediary metabolites from different pathways. This would provide further insights into the metabolic functioning and regulation in *B. methanolicus*. Our results demonstrate the potential of *B. methanolicus* as a versatile platform for the biotechnological production of value-added chemicals from methanol.

## Materials and methods

### Random mutagenesis

Cultures of *B. methanolicus* were grown in MVcMY media containing 6.5 g/L (200 mM) methanol (Supplementary Table 3-5, Additional File 1) at 50°C and 180 rpm for 8-10 h to late exponential phase. Cells were harvested by centrifugation at 4000 x g and the cell pellet was resuspended in 0.1 M sterile phosphate buffer at pH 7. Cultures were then agitated for 5-90 minutes at 50°C in the dark or irradiated with ultraviolet (UV) at 254 nm for 0.5-30 minutes in a shallow petri dish with a constant agitation at room temperature. The mutagen was removed by two washes in 0.1 M phosphate buffer. Cell pellets were ultimately resuspended in a 20% glycerol solution in MVcMY medium and plated onto SOB medium agar plates (44) containing 10 g/L mannitol as serial dilutions for kill rate and colony-forming units (CFU) determination. The mutagenesis conditions resulting in a <99% kill rate (Supplementary Figure 1, Additional File 1) were chosen for screening and selected mutants were stored at -20°C.

### High-throughput mutant screening

Liquid MVcMY medium was inoculated with frozen mutagenized cultures at 0.5% (v/v) inoculum size and grown at 50°C and 180 rpm for 8-10 h. Cultures were then serially diluted and streaked to single colonies onto SOB plates containing 10 g/L mannitol and incubated at 50°C for 48 h. Single colonies were picked with sterile toothpicks into 48-deep-well plates containing 0.7 mL of MvcMY media and incubated at 50°C and 180 rpm until an OD600nm of 0.7-2.5. The seed cultures were then used to inoculate 0.6 mL of ABBM-PM10 production media (Supplementary Table 3-5, Additional File 1) with a 10% (v/v) inoculum size and incubated for 24 h at 50°C and 180 rpm. For GABA production, the cultures were induced with xylose to 10 g/L final concentration at 4 hours into cultivation and acidified to pH 4.6 with 1 M HCl after 24 h of cultivation, then incubated for a further 24 h period. Following incubation, cultures were adjusted to pH 2 with 1 M HCl to lyse cells and release any intracellular product, then centrifuged at 20000 x g for 10 min.

### Molecular cloning

*E. coli* DH5α competent cells were prepared according to the calcium chloride protocol (45). All standard molecular cloning procedures were carried out as described previously (46), or according to manuals provided by manufacturer. The *gad*^St^ gene was amplified using CloneAmp HiFi PCR Premix (Takara) and purified using a QIAquick PCR Purification kit from Qiagen. pBV2xp-*gad*^St^ was constructed by amplifying the *gad*^St^ gene from plasmid pTH1mp-*gad*^St^ with primers PS56 and PS57 (Supplementary Table **Error! Reference source not found.**6, Additional File 1) and joining the resulting PCR product with the *Sac*I and *BamH*I digested pBV2xp by means of the isothermal DNA assembly method (47). The plasmid pBV2xp is a derivative of pHCMC04 for gene expression under control of the *Bacillus megaterium*-derived *xylA* (xylose inducible) promoter (13). Colony PCR were performed using GoTaq DNA Polymerase (Promega) using primers PXPF and BVXR (Supplementary Table **Error! Reference source not found.**6, Additional File 1). The sequence correctness of cloned DNA fragments was confirmed by Sanger sequencing (Eurofins Genomics). *B. methanolicus* MGA3 was made electrocompetent and transformed by electroporation as described by Jakobsen et al (2006) (48). Recombinant *B. methanolicus* strains containing the pBV2xp plasmid were cultivated with the supplementation of 20 µg/mL kanamycin.

### Assessing l-glutamate and GABA production in fed-batch methanol fermentations

For bioreactor experiments testing our newly engineered strains, a two-stage seed strategy was employed. First-stage seed cultures were first inoculated from -20°C working cell banks stored in 20% glycerol into shake flasks with liquid SOB media containing 10 g/L mannitol and incubated for 8-10 h at 50°C and 180 rpm. Shake flasks with ABBM-PM-BR media and without antifoam were inoculated with the second-stage seed with a 10% inoculum size and incubated to an OD600nm of 2-2.5. The cultures were then used to inoculate the bioreactor pre-filled with 2 L of ABBM-PM-BR (Supplementary Table 3-5, Additional File 1) and pre-warmed to 50°C via sterile tubing with a 10% inoculum size. Methanol concentration in the bioreactor was maintained between 4.0-6.1 g/L by automatic feeding of methanol. This was orchestrated via a peristaltic pump controlled by an Arduino microcontroller. The microcontroller was connected to a silicone tube methanol sensor submerged into the fermentation vessel (49). Culture pH was maintained by an ammonia solution (20%), which also served as the nitrogen source. Dissolved oxygen was kept at 30% by supplementation of pure oxygen on demand. Reactors were regularly sampled for OD600nm, pH and ion content (PO ^3-^, SO_4_ ^2-^, NH ^+^), and for l-glutamate and GABA measurements using the same extraction procedure as described above in the “Materials and methods, High-throughput mutant screening” section. To produce GABA, xylose was added to 10 g/L final concentration at 30-40 h post-inoculation. To lower the medium pH, the ammonia feed was shut off until pH 4.5 was reached or the culture was acidified with HCl or H2SO4.

### Batch methanol fermentations for multi-omics sampling

*B. methanolicus* strains ABBM4307, ABBM4307*gad* and MGA3*gad* were cultivated in biological triplicates in 3 L Applikon bioreactors (Getinge, USA) filled with 0.75 L of the multi-omics batch fermentation medium (Supplementary Table 7, Additional File 1). An aeration rate of 0.67 VVM was used and dissolved oxygen (DO) was maintained above 30% throughout the fermentation. The culture temperature and pH were controlled at 50°C and 6.8, respectively. The pH was controlled by the automatic addition of 1 M KOH solution upon demand. The seed cultures were prepared in 2 L baffled glass shake flasks using 400 mL MVcMY medium. Flasks were incubated at 50°C with agitation at 150 rpm until OD600nm of 1.5-2.5 before cells were harvested by centrifugation for 10 min at 3200 x g and 40°C. The cells were resuspended in a small volume of pre-warmed (50°C) batch methanol fermentation medium and the concentrated cell suspension was used to inoculate the bioreactors (1.3%) to a start OD600nm of 0.15. All strains were induced at OD600nm of ca 0.6 by adding xylose to 10 g/L final concentration. The metabolite labelling experiment was initiated by adding 3.2 g/L ^13^C-methanol (99% ^13^C; Eurisotop, France) at OD600nm of 1.6-2.2.

### Multi-omics sampling

Samples for RNA-seq were collected from batch fermentation cultures before and after xylose induction, at approximately OD600nm of 0.5-0.7 and 1.0-1.3, respectively. At the latter RNA-seq sampling point, samples for genomic DNA sequencing were also harvested. For genome sequencing, 50 mL falcon tubes were filled with 20 mL of culture broth, the tubes were centrifuged for 10 min at 8000 x g and 25°C, and cell pellets were stored at -20°C. For RNA-seq sampling, 50 mL falcon tubes were pre-filled with ice and stored at -80°C for 16 h prior to use, to which 20 mL of culture broth samples were immediately added. The tubes were centrifuged for 5 min at 8000 x g and 4°C, and the cell pellets were snap frozen in liquid nitrogen and stored at -80°C.

Supernatant samples were taken to analyze substrate consumption as well as product formation using Nuclear Magnetic Resonance (NMR) measurements. Culture samples of 1.5 mL were taken twelve times throughout the batch cultivation followed by centrifugation for 5 min at 18000 x g and 4°C. Supernatant was then collected and stored at -80°C.

Ion chromatography tandem mass spectrometry (IC-MS/MS) quantification was used to analyze the isotopologues of each metabolite as described previously (50). To determine metabolic pool sizes, blank control samples were taken prior to inoculation of the bioreactors. Later, directly before adding labelled ^13^C-methanol at culture OD600nm of 1.4-1.7, metabolome samples from whole broth (intra- and extracellular pools; WB) and culture filtrate (extracellular pools; CF) were collected using the described previously optimized method (24). CF samples were taken from the bioreactor using a cold 3 mL syringe holding 21 pre-cooled (−20°C) stainless steel beads, attached to a 0.20 μm filter (Whatman Puradisc). WB samples were collected with the same approach, but without the filter. Samples were transferred to ice cold 1.5 mL tubes from which technical triplicates of 150 µL were dispensed into new 5 mL extraction tubes containing 3 mL extraction solution (acetonitrile:methanol:0.1 M formic acid in 40:40:20 (v/v/v)) pre-chilled to -20°C prior to use. Samples were homogenized by vortexing. To each tube (except for blank control samples), 200 µL of internal standard was added (cell extract of *E. coli* cultivated on 99% [U-^13^C6]-glucose, Eurisotop, France). The extracted samples were stored at −80°C. The average time between sample collection and extraction of technical triplicates was 18 ± 2 sec for WB samples and 25 ± 3 sec for CF samples.

Label enrichments in the intracellular metabolites were sampled directly after performing the addition of 3.2 g/L ^13^C-methanol. WB and CF samples were collected as described above but using cold 1 mL syringes with 7 stainless steel beads, and without adding any internal standard. From each bioreactor, technical duplicates of 13 WB and 3 CF samples were collected within 3.5 min average. Samples were stored at -80°C.

The dry cell weight (DCW) was determined by sampling 20 mL of culture from all bioreactors when OD600nm reached 2.5-3.0. The samples were centrifuged for 10 min at 3200 x g and 4°C, pellets thoroughly washed and eventually transferred to pre-weighted aluminum cups. The samples were dried in an oven at 105°C for 24 h, along with blank controls. A conversion factor of 0.24 g/L DCW per OD600nm unit was obtained.

### Analytical procedures

In the fed-batch fermentations, sampled supernatants were routinely analyzed for l-glutamate and GABA content by high performance liquid chromatography (HPLC) in a AkzoNobel Kromasil 100-5-C18 200 x 4.6 column, using 0.1% HFBA in a water:methanol mixture (80:20 (v/v)) as the mobile phase.

Metabolite concentration in culture supernatants used in MFA were measured by 1D 1H NMR at 280K using a 30° angle pulse 16 scans, and a pre-saturation of the water signal (zgpr30) was applied during a relaxation delay of 3s (total repetition time of 7s) in an Avance III 800 MHz spectrometer (Bruker, Germany) equipped with a QCI-P 5mm cryoprobe (Bruker, Germany). Deuterated trimethylsilyl propionate (TSP-d4) dissolved in D2O was used as an internal standard for quantification. The spectra were processed using Topspin 3.5. Metabolite pool sizes were quantified by (IC-MS/MS) using the U-^13^C-labeled *E. coli* cell extract as an internal standard (51).

Metabolite quantification for fluxomics analysis was performed with liquid anion-exchange chromatography as described previously (50). In short, high-resolution experiments were performed with an Ultimate 3000 high-performance liquid chromatography (HPLC) system (Dionex, CA, USA) coupled to an LTQ Orbitrap Velos mass spectrometer (Thermo Fisher Scientific, Waltham, MA, USA) equipped with a heated electrospray ionization probe. MS analyses were performed in positive Fourier transform MS (FTMS) mode at a resolution of 60000 (at 400 m/z) in full-scan mode, with the following source parameters: capillary temperature, 275°C; source heater temperature, 250°C; sheath gas flow rate, 45 AU (arbitrary unit); auxiliary gas flow rate, 20 AU; S-lens radio frequency (RF) level, 40%; source voltage, 5 kV. Isotopic clusters were determined by extracting the exact mass of all isotopologues, with a tolerance of 5 ppm. The following metabolites analyzed were used in MFA analysis: fructose 1,6-bisphosphate (FruBP), AKG, Mal, Cit, 3-phosphoglycerate (3PG) plus 2-phosphoglycerate (PGA), ribose 5-phosphate (Rib5P) plus Ribu5P, plus xylose 5-phosphate (Xyl5P), glucose 6-phosphate (Glc6P), Fru6P, fumarate (Fum), phosphoenolpyruvate (PEP) and aconitate (Aco). After peak integration, the raw peak areas were corrected for the contribution of naturally abundant isotopes using the IsoCor software version v2.1.3 (52).

### Calculation of growth, substrate uptake and degradation rates

Quantitative cell growth parameters, extracellular flux and evaporation constant from the measured time-dependent concentrations of biomass and extracellular metabolites measured by NMR were inferred using Physiofit version v3.3.6 (53). A conversion factor was used to obtain cellular DCW from OD600nm units (0.24 gDCW/OD600nm) and calculate specific substrate and production rates. Averages and standard deviations were calculated based on three biological replicates.

### Variant Calling

Genomic DNA was isolated using the NucleoSpin® Microbial DNA kit (Macherey-Nagel). Long and short DNA reads were generated by Nanopore and Illumina sequencing, respectively. For library preparation, the TruSeq DNA PCR-free high-throughput library prep kit (Illumina) and the SQK-LSK109 sequencing kit (Oxford Nanopore Technologies [ONT]) were used without prior shearing of the DNA. To generate the short reads, a 2 × 300-nucleotide run (MiSeq reagent kit v3, 600 cycles) was executed. The long reads were generated on a GridION platform using a R9.4.1 flow cell. Base calling and demultiplexing were performed using GUPPY v3.2.10 with the high accuracy base calling model. Assemblies were done using CANU v1.8 (54) for the Nanopore long read data, followed by polishing with medaka v.1.0.1 (https://github.com/nanoporetech/medaka) and pilon v1.22 (55). The polished genomes were then used to identify SNPs with snippy v.4.0-dev2 (https://github.com/tseemann/snippy) in contig mode. To interpret the effect of SNPs in the reaction sites of altered proteins, their amino acid sequences were submitted to functional analysis in the InterProScan software (56).

### RNA-seq analysis

For the total RNA extraction from the tested *B. methanolicus* strains, we employed the NucleoSpin RNA isolation kit (Macherey-Nagel) and the RNase-free DNase set (Qiagen), following the manufacturers’ protocols. Snap frozen bacterial pellets were thawed on ice, and total RNA isolation was carried out individually for each replicate of the cultivation condition. To ensure the quality of the resulting RNA material and confirm the absence of DNA contamination, PCR was performed employing gene-specific forward primer KATFW and reverse primer KATREV (Supplementary Table **Error! Reference source not found.**6, Additional File 1). They bind specifically to the catalase encoding gene *katA* in *B. methanolicus*. Furthermore, the concentration and quality of the samples were assessed by measuring the 260/280 nm and 260/230 nm ratios using a NanoDrop^TM^ One (Thermo Scientific^TM^). The RNA quality was checked by Trinean Xpose (Trinean NV, Gentbrugge, Belgium) and Agilent RNA 6000 Nano Kit with an Agilent 2100 Bioanalyzer (Agilent Technologies, Böblingen, Germany). rRNA was removed from total RNA with Ribo-Zero rRNA Removal Kit (Illumina, San Diego, CA, USA), and removal of rRNA was checked using Agilent RNA 6000 Pico Kit and an Agilent 2100 Bioanalyzer (Agilent Technologies). cDNA libraries were prepared with TruSeq Stranded mRNA Library Prep Kit (Illumina), and the resulting cDNA was paired-end sequenced using an Illumina NextSeq 500 system with a 2 × 75 bp read length.

Paired-end reads generated from the RNA-seq were aligned to the *B. methanolicus* MGA3 genome reference sequences, encompassing the chromosome as well as the two plasmids pBM19 and pBM69 (GenBank accession numbers CP007739, CP007741, and CP007740, respectively). Before the alignment, the reads underwent trimming with a minimum length of 36 base pairs using the Trimmomatic v0.33 tool (57). The trimmed reads were then aligned to the reference sequences using the short-read alignment software Bowtie 2 (58). The ReadXplorer software v2 (59) was employed for visualizing the mapped reads, conducting operon structure analysis, and determining reads per kilobase per million mapped reads (RPKM) values. Furthermore, the differential gene expression analysis was executed with the statistical method DESeq2 (60) within the same software framework. To classify a gene as differentially expressed, the criteria included a change in expression level exceeding 50, with the adjusted *p*-value set at or below 0.01. For differentially expressed genes (DEGs) encoding proteins of unknown function, BLASTx analysis (61) was applied to identify conserved protein family. Venn diagrams were generated using InteractiVenn software (62).

### Metabolic flux analysis model and simulations

A model previously designed that covers the central carbon metabolism and biomass needs of *B. methanolicus* MGA3 (15) was modified to adjust to the present study, i.e., network topology changes to include l-glutamate production and the GABA synthetic pathway. The influx_i uses a nonstandard legacy file format (FTBL) to encode the metabolic network and the associated atom-atom transitions. This format centralizes the metabolic network with all biological measurements, i.e., metabolite pool sizes, fluxes, and carbon isotopologue distributions. Simulations needed for non-stationary ^13^C-MFA were performed with influx_si v5.3.0 in non-stationary mode (63). The influx software has the advantage of allowing for the integration of labelling data coming from different experimental setups (MS and NMR) and even supporting several integration strategies (stationary, nonstationary, and parallel labelling).

Experimental data were fitted to our models as described above. For each culture replicate, a chi-squared goodness-of-fit statistical test was performed to ensure that simulated data for each biological replicate was significantly close to experimental data. All tests were significant with a significance level (α) of 0.05.

### Genome-scale model analysis

To reconstruct the GSM of *B. methanolicus*, a bioinformatic pipeline described by Peyraud et al., 2016 was used (64). In this approach, orthologs, i.e., genes which are likely to perform the same function, are identified between a query genome and a reference genome for which there is already a GSM available. Then the gene protein reaction (GPR) is extracted from the reference GSM and combined with ortholog data allowing the propagation of GPR toward the query genome. Genomes of two microorganisms (*E. coli* and *B. subtills*) for which GSM were available were used as references. *E. coli* was selected because its GSM is well annotated. The “iAF1260” model of *E. coli* (65) containing 2382 reactions for 1972 metabolites was chosen as the first reference GSM for the reconstruction. However, *E. coli,* being a Gram-negative bacterium, does not have the membrane specificities associated with Gram-positive bacteria, and for that reason *B. subtilis* was chosen as the second reference. The “iBsu1103” GSM was used, which includes 1681 reactions for 1625 metabolites (66). Since these metabolic models use different ontologies, the identifiers were standardized using a tool called SAMIR (Semi Automatic Metabolic Identifier Reconciliation) (64).

The orthology prediction being based on protein sequence in order to avoid codon bias, proteins sequences of the entire genome (i.e., proteome) of the reference and source organisms were download from UNIPROT (67). The orthology prediction was made using a tuned version of Inparanoid (68). Inparanoid found orthologs between the proteome of *B. methanolicus* and each of the two reference proteomes. As a result, out of the 3301 protein sequences of the *B. methanolicus* proteome, 2131 orthologues were detected against the proteome of *E. coli* (containing 10530 sequences) and 2154 against that of *B. subtilis* (containing 8308 sequences). Next, the Autograph method (69) was used to automatically transfer by orthology the GPRs of the two reference GSM in one draft model. In this step, each query gene for which there is an orthology with the reference gene(s) is associated with the corresponding reaction(s) from the reference model. By combining the orthology data with the GPRs from the reference GSM, 764 reactions were propagated from “iBsu1103” and 1642 from “iAF1260”. Among these reactions, reactions with complete (918), incomplete (806) or without (682) GPR can be distinguished. The so-called full GPR reactions are reactions where all the genes involved in a reaction in the reference model have been associated with genes present in *B. methanolicus*. Conversely, reactions without GPR are reactions for which no ortholog has been identified between the reference and the target. These reactions are still propagated to avoid losing information. 87.1% of these reactions were spontaneous exchange and transport reactions that are not encoded by genes. Finally, the so-called incomplete GPR reactions are those for which all the genes attributed to a reaction in the reference model have not been associated with genes *in B. methanolicus*. These GPRs relate to reactions catalyzed by enzymatic complexes or isoenzymes. At this stage the draft reconstruction enclosed 1642 and 764 propagated from *E. coli* and *B. subtillis*, respectively. Finally, a manual refinement was performed. First, the draft reconstruction was cleaned by removing the redundant GRPs, the non-functional reactions with incomplete or no GPRs, the dubious homology and the periplasmic transport and exchange reactions (originating from the *E. coli* model). Unlike *E. coli*, *B. methanolicus* does not possess a periplasm and therefore no periplasmic exchange and transport reactions either. Then reconstruction was further improved by adding specific GPRs (e.g., methylotrophic metabolism, biomass reactions). The biomass reactions were adapted from those in the model “iBsu1103”. Metexplore (70) was used to identify the “dead ends” (i.e., metabolites which are produced but not consume or inversely) and visualize the metabolic pathways. MetaCyc (71) was used to identify the reactions and Blastp (https://blast.ncbi.nlm.nih.gov/Blast.cgi?PAGE=Proteins) to associate these with the corresponding genes. Reactions and metabolites identifiers were updated to the latest BiGG identifier and genes were associated with a RefSeq number. The “iBmeth613” model of *B. methanolicus* contains 1020 reactions, 899 metabolites, 613 genes and 2 compartments (cytosol and extracellular space). The quality of this model was tested by performing a mass balance analysis of all the reactions except biomass and “MAGIC” reactions that synthesize abstract metabolites for the biomass.

The functions “parsimonious flux balance analysis” (pFBA) and “flux variability analysis” (FVA) of the COBRA toolbox for Python version 3.8.8 (72) and the “iBmeth613” model of *B. methanolicus* (this study) were used to simulate the flux distribution and flux changes. The objective function was the growth rate, and the model was constrained with the uptake and production rates (in mmol/gDCW/h) calculated from the MFA analysis for MGA3*gad* and ABBM4307*gad* (Supplementary Table 1, Additional File 3). pFBA aims to identify a realistic flux distribution within a model by minimizing the squared sum of all fluxes while preserving the achieved optimum. FVA, on the other hand, assesses the minimum and maximum bounds for each reaction flux that can still meet the constraints, employing a maximization followed by a minimization for each reaction in the GSM.

## Declarations

### Ethics approval and consent to participate

Does not apply

### Consent for publication

Does not apply

### Availability of data and materials

The datasets supporting the conclusions of this article are included within the article and its additional files. Genomic data is available under BioProject PRJNA1167228. Transcriptome data was deposited in the Gene Expression Omnibus (GEO) database (GSE277855). MFA and GSM raw models, input and output files and statistical analysis results are available in FAIRDOM (https://fairdomhub.org/data_files/7513?version=1 and https://fairdomhub.org/data_files/7516?version=1).

### Author’s contributions

DG, MI and DV engineered the strains used in this study. LFB, MI, and IN conducted the multiomics fermentation and sampling; TBu, CH, and CR performed the sequencing work and variant calling; LFB carried out the RNA-seq analysis; CMV performed the ^13^C-labeling data analysis and MFA simulations; SH developed the GSM and performed the GSM simulations. LFB, SH, MI, IN, CMV and DV drafted the original manuscript. LFB and CMV edited and prepared the final version of the manuscript. TBr, SH, IN, GK, and VFW secured the funding necessary for the completion of this study. All authors reviewed the manuscript.

## Acknowledgements

We thank MetaToul-FluxoMet (Toulouse, France), which is part of the French National Infrastructure for Metabolomics and Fluxomics (https://www.metabohub.fr) and funded by the ANR (MetaboHUB-ANR-11-INBS-0010), for their help in collecting, processing, and interpreting ^13^C NMR and MS data. We also thank Pierre Millard (INRAE, Toulouse, France) for insightful discussions on MFA, as well as Baudoin Delépine for his help with the first draft of the GSM. We would also like to thank the support of the entire Acies Bio lab team, especially Petra Kralj with the random mutagenesis and screening, Dr. Jaka Horvat with the analytics and processing of samples, as well as Toni Nagode and Filips Oleskovs with the bioreactor fermentations.

