## Additional File 1 for "Systems-level analysis provides insights on methanol-based production of L-glutamate and its decarboxylation product γ-aminobutyric acid by *Bacillus methanolicus*"

#### Table of contents

Supplementary Figures

*Supplementary Figure 1. Schematic representation of high-throughput mutagenesis and screening protocol.*
*Supplementary Figure 2. Correlation between estimated (measured by MFA) and predicted (simulated with GSM).*
*Supplementary Figure 3. Schematic representation of the aerobic electron transport chain in* B. methanolicus.

Supplementary Tables

*Supplementary Table 1. Summary of sequencing reads and mapping statistics for whole transcriptome libraries derived from* B. methanolicus *RNA samples.*
*Supplementary Table 2. Differentially expressed gene (DEG) counts in RNA-seq libraries of four* B. methanolicus *strains MGA3*gad*, ABBM4307, ABBM4307*gad *and ABBM4307*gad *NI.*
*Supplementary Table 3. Composition of the media used during classical mutagenesis and fed-batch cultivations.*
*Supplementary Table 4. Composition of MVcM trace metals solution (1000x).*
*Supplementary Table 5. Composition of the MVcM complete vitamins solution (1000x).*
*Supplementary Table 6. Primers used in this study.*
*Supplementary Table 7. Composition of the batch methanol fermentation medium used for multi-omics cultivations and sampling.*

#### Supplementary Figures

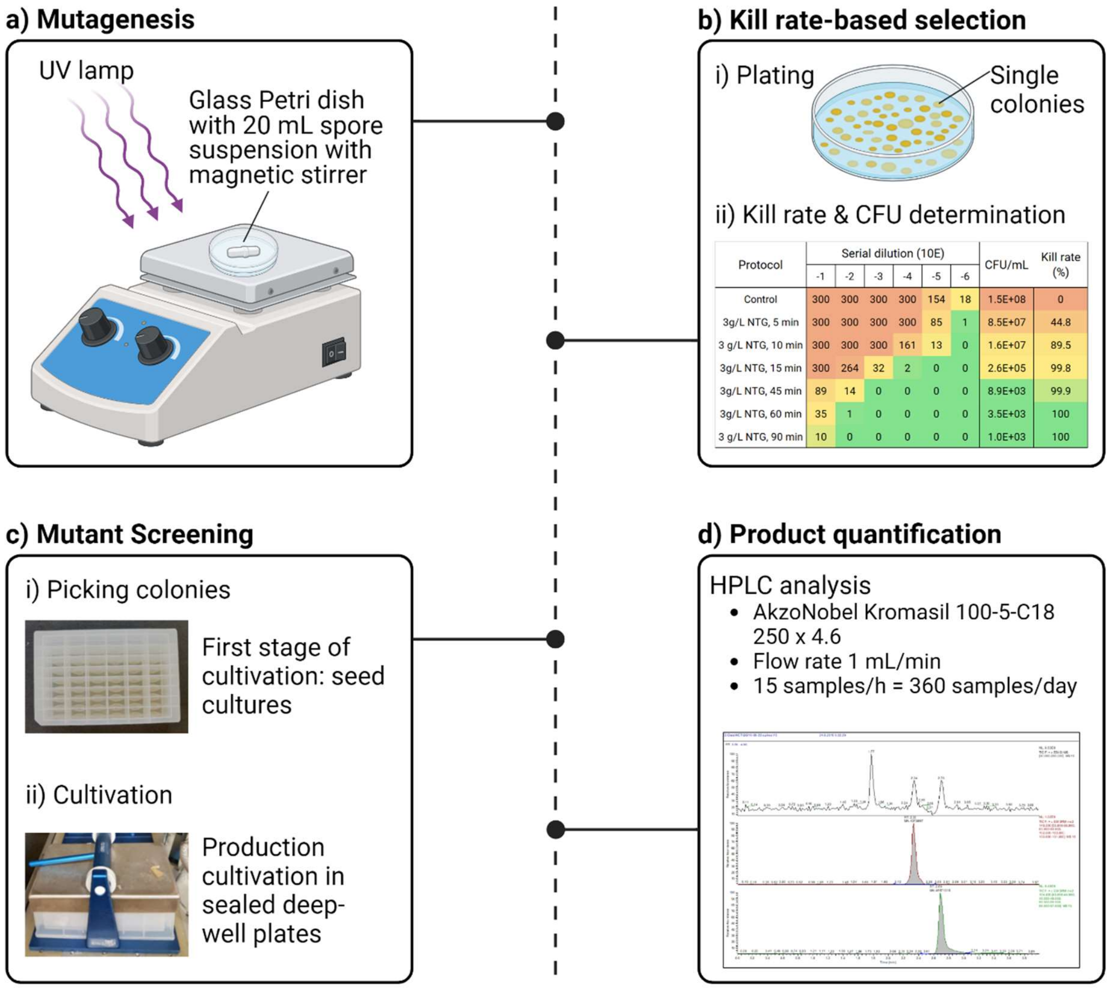


*Supplementary Figure 1. Schematic representation of high-throughput mutagenesis and screening protocol.*

*a) Cultures were mutagenized with NTG, EMS or UV (exemplified here) under a range of conditions (time, concentration, UV lamp distance). b) After mutagenesis, cultures were serially diluted and plated. For each mutagenesis, the condition that resulted in a <99% kill rate was chosen for screening. c) Single colonies were picked into 48-well plates containing SOB liquid media and grown overnight. The overnight cultures were then inoculated into ABBM-PM10 production media containing both methanol and mannitol, as well as MOPS buffer. d) Cultures were sampled after 24 h and the L-glutamate content of the supernatant was determined by HPLC with a throughput of 360 mutants per day. Figure generated in the BioRender platform (BioRender.com/k43l501).*

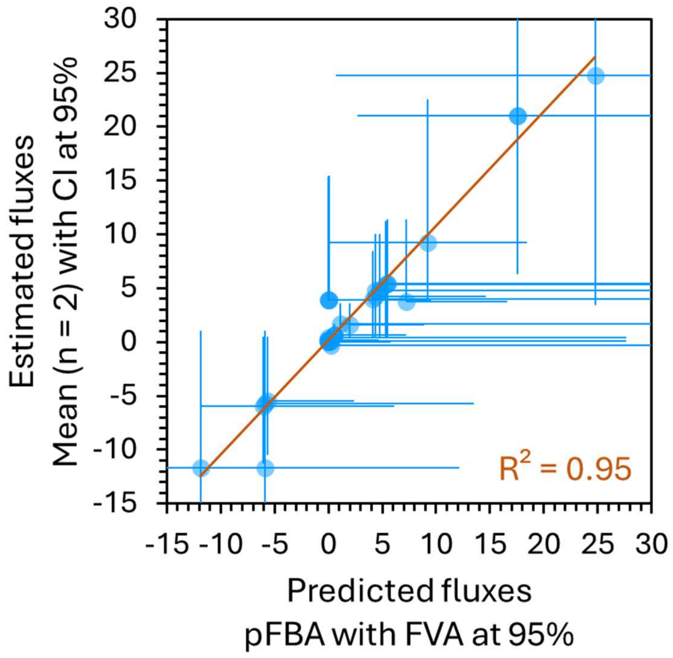


*Supplementary Figure 2. Correlation between estimated (measured by MFA) and predicted (simulated with GSM).*

*By constraining the input and output with the estimated values (from $^{13}C$ experiment), the GSM optimization (made on the biomass reaction) can find the optimal estimated reaction rate with a strong correlation (correlation coefficient between predicted and estimated values is ~0.95). See the "Materials and methods" section for more details.*

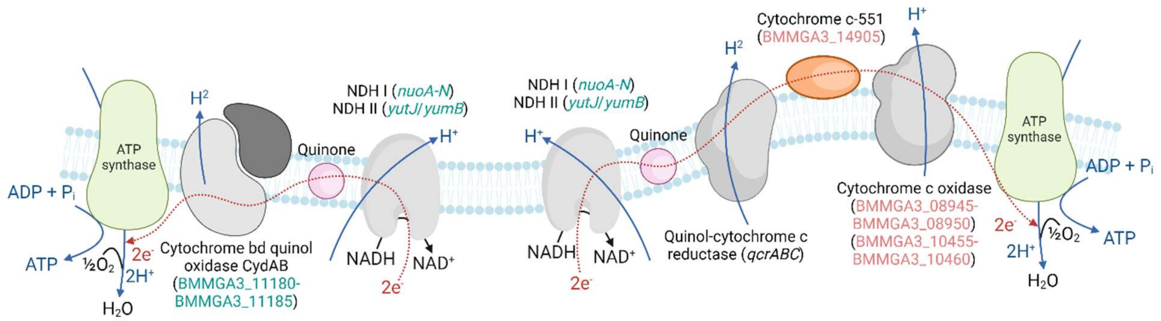


*Supplementary Figure 3. Schematic representation of the aerobic electron transport chain in* B. methanolicus.

*Genes shown in green are upregulated, while those in red are downregulated in the ABBM4307*gad *strain compared to MGA3*gad *based on transcriptomic analysis. NDH- NAHD-quinone oxidoreductase. Figure generated the in BioRender platform (BioRender.com/z23j212).*

### Supplementary Tables

*Supplementary Table 1. Summary of sequencing reads and mapping statistics for whole transcriptome libraries derived from* B. methanolicus *RNA samples.*

*The results of number of paired reads after trimming (minimal length of 36 nucleotides) and overall alignment rate after mapping are represented as average and standard deviation of technical triplicates.*

| Strain/condition | Reference | Paired reads (n) | | | Alignment (%) | | |
|---|---|---|---|---|---|---|---|
| MGA3*gad* | Chromosome | 1,213,269 | ± | 52,948 | 89.0 | ± | 0.5 |
| | pBM19 | | | | 6.4 | ± | 0.2 |
| | pBM69 | | | | 1.0 | ± | 0.1 |
| | pBV2xp-*gad*$^{St}$ | | | | 3.2 | ± | 0.3 |
| ABBM4307 | Chromosome | 1,184,180 | ± | 91,729 | 93.2 | ± | 0.8 |
| | pBM19 | | | | 5.8 | ± | 0.7 |
| | pBM69 | | | | 0.7 | ± | 0.1 |
| | pBV2xp-*gad*$^{St}$ | | | | 0.0 | ± | 0.0 |
| ABBM4307*gad* | Chromosome | 1,281,976 | ± | 41,888 | 92.2 | ± | 0.1 |
| | pBM19 | | | | 6.1 | ± | 0.1 |
| | pBM69 | | | | 0.7 | ± | 0.0 |
| | pBV2xp-*gad*$^{St}$ | | | | 0.6 | ± | 0.0 |
| ABBM4307*gad* NI | Chromosome | 1,102,126 | ± | 52,128 | 89.3 | ± | 0.3 |
| | pBM19 | | | | 6.2 | ± | 0.1 |
| | pBM69 | | | | 0.8 | ± | 0.0 |
| | pBV2xp-*gad*$^{St}$ | | | | 3.4 | ± | 0.1 |

*Supplementary Table 2. Differentially expressed gene (DEG) counts in RNA-seq libraries of four* B. methanolicus *strains MGA3*gad*, ABBM4307, ABBM4307gad and ABBM4307*gad *NI.*

| DEG counts | Chromosome | | pBM19 | | pBM69 | | pBV2xp-*gad*[St] | |
|---|---|---|---|---|---|---|---|---|
| | **Down** | **Up** | **Down** | **Up** | **Down** | **Up** | **Down** | **Up** |
| ABBM4307*gad* vs MGA3*gad* | 421 | 553 | 6 | 7 | 11 | 14 | 1 | 3 |
| ABBM4307*gad* vs ABBM4307*gad* NI | 206 | 194 | 1 | 1 | 4 | 2 | 4 | 4 |
| ABBM4307*gad* vs ABBM4307 | 17 | 31 | 10 | 16 | 17 | 33 | n.d. | n.d. |

*Supplementary Table 3. Composition of the media used during classical mutagenesis and fed-batch cultivations.*

| **Component** | **MVcMY** | **ABBM-PM10** | **ABBM-PM-BR** |
|---|---|---|---|
| $K_2HPO_4$ | 4.1 g/L | 4.1 g/L | 6.2 g/L |
| $NaH_2PO_4$ | 1.3 g/L | 1.3 g/L | 2.0 g/L |
| $(NH_4)_2SO_4$ | 2.1 g/L | 4.0 g/L | 4.0 g/L |
| Mannitol | / | 5.0 g/L | / |
| MOPS | / | 20.9 g/L | / |
| Yeast extract | 0.25 g/L | / | 1 |
| 1M $MgSO_4 \cdot 7H_2O$ | 1 mL/L | 0.2 mL/L | 0.2 mL/L |
| MVcM complete vitamins 1000X | 1 mL/L | 0.2 mL/L | 8 mL/L |
| MVcM trace metals 1000X | 1 mL/L | 1 mL/L | 1 mL/L |
| Biotin | / | / | 12 mg/L |
| Antifoam SAG | / | / | 1 mL/L |
| Methanol | 6.5 g/L | 6.5 g/L | 5.6 g/L |

*Supplementary Table 4. Composition of MVcM trace metals solution (1000x).*

| Component | g/L |
|---|---|
| $FeSO_4 \cdot 7H_2O$ | 5.56 |
| $CuCl_2 \cdot 2H_2O$ | 0.027 |
| $CaCl_2 \cdot 2H_2O$ | 7.35 |
| $CoCl_2 \cdot 6H_2O$ | 0.040 |
| $MnCl_2 \cdot 4H_2O$ | 9.90 |
| $ZnSO_4 \cdot 7H_2O$ | 0.288 |
| $Na_2MoO_4 \cdot 2H_2O$ | 0.048 |
| $H_3BO_3$ | 0.031 |
| Concentrated HCl | 72.72 |
| $ddH_2O$ | up to 1L |

*Supplementary Table 5. Composition of the MVcM complete vitamins solution (1000x).*

| Component | Amounts (g/L) |
|---|---|
| d-Biotin | 0.100 |
| Thiamine·HCl (vitamin B1) | 0.100 |
| Riboflavin (vitamin B2) | 0.100 |
| Pyridoxine·HCl | 0.100 |
| Pantothenate | 0.100 |
| Nicotinamide (vitamin B3) | 0.100 |
| p-Aminobenzoic acid (vitamin L1) | 0.020 |
| Folic acid (vitamin B11) | 0.010 |
| Alphamine (vitamin B12) | 0.010 |
| Lipoic acid (thioctic acid) | 0.010 |
| dd$H_2O$ | up to 1L |

*Supplementary Table 6. Primers used in this study.*

| Name | Sequence 5'->3' | Function |
|---|---|---|
| PS56 | TTCACTTAAGGGGGAAATGGCAAatgAAAAAAGCA TCGGAGGAAGT | *gad*$^{St}$, FW primer |
| PS57 | ACGACGGCCAGTGAATTCGAGCTtcaATGATGAAA CGGTTTATGTTGG | *gad*$^{St}$, RV primer |
| PXPF | TGTTTATCCACCGAACTAAG | sequencing primer for pBV2xp; FW |
| BVXR | CCGCACAGATGCGTAAGGAG | sequencing primer for pBV2xp; RV |
| KATFW | GCAAATAACTACCAGCGTGA | *katA*, FW primer |
| KATRV | TCACGCTGGTAGTTATTTGC | *katA*, RV primer |

*Supplementary Table 7. Composition of the batch methanol fermentation medium used for multi-omics cultivations and sampling.*

| **Component** | **Amounts** |
|---|---|
| $Na_2HPO_4$ | 1.38 g/L |
| $KH_2PO_4$ | 0.61 g/L |
| $NH_4Cl$ | 2.50 g/L |
| Yeast extract | 0.048 g/L |
| 1 M $MgSO_4{}^{-}7H_2O$ | 1 mL/L |
| MVcM complete vitamins 1000X | 1 mL/L |
| MVcM trace metals 1000X | 1 mL/L |
| Methanol | 4.8 g/L (150 mM) |
